# Activity-dependent organization of prefrontal hub-networks for associative learning and signal transformation

**DOI:** 10.1101/2021.08.31.458461

**Authors:** Masakazu Agetsuma, Issei Sato, Yasuhiro R Tanaka, Luis Carrillo-Reid, Atsushi Kasai, Yoshiyuki Arai, Miki Yoshitomo, Takashi Inagaki, Hitoshi Hashimoto, Junichi Nabekura, Takeharu Nagai

## Abstract

Associative learning is crucial for adapting to environmental changes. The encoding of associative learning involves the dorso-medial prefrontal cortex (dmPFC), and is underpinned by interactions within the resident neuronal population. However, the nature of this population coding is poorly understood. Here we developed a pipeline for computational dissection and longitudinal two-photon imaging of neural population activities in the mouse dmPFC during fear-conditioning procedures, enabling us to detect learning-dependent changes in the dmPFC topology. Through regularized regression methods and graphical modeling, we found fear conditioning organized neuronal ensembles encoding conditioned responses (CR), with enhancing their coactivity, functional connectivity, and association with conditioned stimuli (CS). This suggests that fear conditioning drives dmPFC reorganization to generate novel associative circuits for CS-to-CR transformation. Importantly, neurons strongly responding to unconditioned stimuli (US) during conditioning anterogradely became a hub of the CR ensemble. Altogether, we demonstrate learning-dependent dynamic modulation of population coding structured on an activity-dependent hub-network formation within the dmPFC.

**Teaser:** Optical and computational dissection uncovered how prefrontal cortical networks are rewired to encode new associative memory

**Significance statement:** Animals learn to adapt to changing environments. Associative learning is one of the simplest types of learning that has been intensively studied over the past century. Recent development in molecular, genetic, and optogenetic methods has enabled the identification of a neural population encoding the associative memory in the brain. However, it remains unclear how information is stored and processed by the neural population to encode and retrieve the associative memory. To investigate the nature of this population coding, we developed an optical and computational dissection method, demonstrating how associative learning drives reorganization of the neural network in the dorso-medial prefrontal cortex and generates novel circuits for associative memory and signal transformation.

## Introduction

Animals learn to adapt to changing environments for survival. Associative learning, such as classical conditioning, is one of the simplest types of learning that has been intensively studied over the past century (*1*, *2*). It is based on repeated pairings of a neutral conditioned stimulus (CS) such as a tone, and an unconditioned stimulus (US) such as foot shock, that eventually makes the subjects respond to the CS by itself and elicit a conditioned response (CR), e.g., freezing behavior in the associative fear learning paradigm. During the last two decades, technical development in molecular, genetic, and optogenetic methods has enabled the identification of a population of neurons in the brain, named the memory engram, which encodes and regulates associative memory (*3*). How information is stored and processed by the neural population to encode and retrieve the associative memory, however, remains unclear (*3*). In addition, although previous studies proposed the possibility that the formation of associative memory may involve novel associative connections between the originally distinct CS and US networks to enable the CS-to-CR transformation, direct evidence is quite limited.

The prefrontal cortex (PFC) is a brain region that regulates associative fear memory, which is evolutionarily conserved in mammals, from humans to primates to rodents (*4*–*9*). Dysfunction of the PFC may lead to various psychiatric diseases including the post-traumatic stress disorder (*10*), and the associative fear memory paradigm has been used as a research model to investigate the underlying mechanisms of this disorder. The dorsal part of the medial prefrontal cortex (dmPFC) of rodents is a brain region demonstrated to be important for the retrieval of associative fear memory (*11*–*16*). During fear memory retrieval and evoked freezing responses (i.e., CR), activated individual neurons (*17*) or an enhanced synchrony of neural populations (*14*) in the dmPFC are observed, while pharmacological or optogenetic silencing of the dmPFC and its projections to specific downstream targets suppresses fear memory retrieval (*11*, *12*), revealing that associative fear memory is normally stored in the dmPFC. Therefore, the dmPFC can serve as an interesting target to address the fundamental question of what structural and computational alterations in the prefrontal networks are required to organize novel associative memories. Also, the study may contribute to our understanding of how novel associative memory is stored in the dmPFC together with pre-existing networks such as those regulating sensory and motor information.

To address these points, here we developed a pipeline for computational dissection and longitudinal imaging of neural population activities in the dmPFC during fear conditioning procedures in mice, which enabled us to uncover learning-dependent changes in the internal topology, functional connectivity and computational architecture of the dmPFC upon memory acquisition.

## Results

### Longitudinal imaging of neural population activities in mouse dmPFC during fear-conditioning procedures

To perform longitudinal imaging of neuronal population activities in the dmPFC during fear-conditioning procedures in mice, we first developed a system to perform cued-fear conditioning and memory retrieval while imaging awake and behaving mice with a two-photon microscope (Fig. 1), which enabled us to record the neural activities of hundreds of neurons with single-cell resolution and to further elucidate experience-dependent changes in functional connectivity in the dmPFC internal network, as shown later. The mice were head-fixed under the microscope objective and placed on a running disk, which was used to record the mouse locomotion status (e.g., locomoting, stationary, or expressing a freezing response) (Fig. 1A). Tones and foot shocks were delivered as the CS and US, respectively. Two different tones were used; one was associated with the US (CS+) and the other was not (CS−), as described in previous studies (*14*, *18*). On day 3 (D3), the mice underwent a habituation session, in which they received 4 presentations of the CS− and CS+ alternately. The habituation session was immediately followed by discriminative fear conditioning session on the same day, in which only the CS+ was paired with the US (Fig. 1B). The US was delivered during the last 1-s of each 30-s CS+ trial. The CS− and CS+ trials were performed alternately (inter-trial intervals, 50–150 s). The next day (D4), the conditioned mice underwent a retrieval and extinction session, in which they received 4 presentations of the CS− and 12 presentations of the CS+ (4 presentations of the CS− and CS+ trials alternately, followed by 8 CS+ trials; Fig. 1B). Behavioral analyses revealed that the mice learned to exhibit freezing-like behavior, i.e., decrease their locomotion as a conditioned response (CR), specifically during the CS+, only after the fear conditioning (Fig. 1B, C). On the other hand, as reported previously (*14*, *18*), the CR observed during the early phase on D4 was extinguished after repeated exposure to the CS+ only (Figs. 1D, E). Overall, these behavioral data established that our behavioral system and the fear conditioning protocol were useful for observing a change in the neural representation during associative fear learning.

**Fig. 1.**
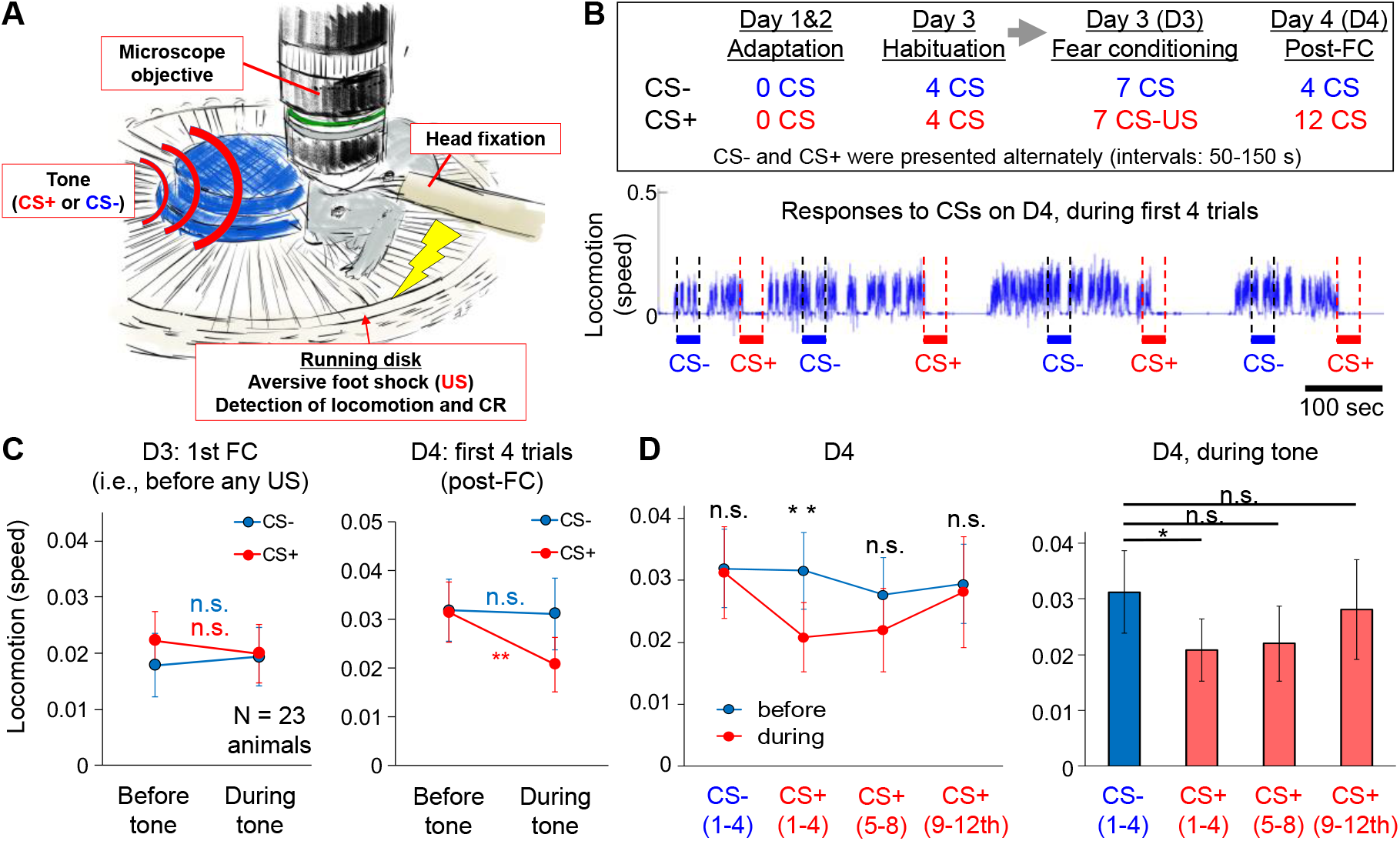
Cued fear conditioning during two-photon microscopy. (**A**) Schematic diagram showing the system used to perform the cued-fear conditioning and memory retrieval under a two-photon microscope. (**B**) (top) Experimental protocol. CS, conditioned stimulus; US, unconditioned stimulus; FC, fear conditioning. (bottom) An example of the changes in locomotion over time of a mouse on day (D) 4 (first four trials). (**C-D**) Fear conditioning under the microscope produced CS+-specific memory consolidation. Comparisons of the locomotor speed between before the tone onset and during the tone presentation are shown in (C-D). Before the fear conditioning (on D3), the mice (N = 23) exhibited no significant change in locomotion during the CS+ and CS− presentations (C, left, and D). After the fear conditioning (i.e., during fear retrieval; the first four trials on D4), however, the CS+ suppressed locomotion as a CR, while the CS− induced no significant change (C, right, and D). After repeated presentations of the CS+ (fear extinction; 5th-12th trials on D4), the CS+-evoked CR became smaller until no significant change in locomotion was observed upon CS+ presentation (D). (**E**) Statistical comparison among responses to the CS− and those to the CS+ at each testing phase on D4 during the tone presentation revealed that locomotion during CS+ was significantly lower only during trials 1–4 on D4, and not after repeated presentations to the CS+ (5th-12th trials). Note that locomotion during pre-tone-onset (before) was not significantly different between the CS− and CS+ conditions. *p<0.05; **p<0.01; n.s., not significant by Wilcoxon signed-rank test (the Friedman test followed by post-hoc multiple comparisons revealed similar results for panel E). Error bars, s.e.m.

Next, to monitor the neural activities in the dmPFC by two-photon microscopy, we implanted a 2-mm microprism along the rostral midline of the brain to optically access the dmPFC region. Although the size of the prism was larger than that of prisms used in previous work (*19*),there was sufficient space and no callosal fibers between the hemispheres around the dmPFC area, especially at the rostral region, enabling the smooth insertion of the prism without cutting prefrontal or callosal neural fibers (Fig. 2A). Using a genetically encoded Ca^2+^ indicator, GCaMP6f, expressed by an adeno-associated virus (AAV), the activities from a wide region of the prefrontal area were chronically visualized (Fig. 2B, C and Movie S1). To specifically record the activity of the excitatory neurons (*20*) and separate them from inhibitory neurons that may have a distinct function in the dmPFC (*13*), the GCaMP6f was expressed under regulation of the CaMKII promoter (*21*, *22*). The CS and US presentation did not disturb image acquisition (Movie S2). We focused on analyzing activities in the dmPFC area (see the Materials and Methods for details). In most of the data analyses, the neural representation during the first 3 trials of the fear conditioning on D3 (D3-early [D3E]) were compared with those during the first 3 trials on D4 (D4-early [D4E]) to assess the changes occurring after the fear conditioning and memory consolidation. The data obtained during the last 3 trials on D3 (D3-late [D3L]) were used to assess the late conditioning phase, and the data obtained during the last 3 trials on D4 (D4-late [D4L]) were used to assess the extinction phase.

**Fig. 2.**
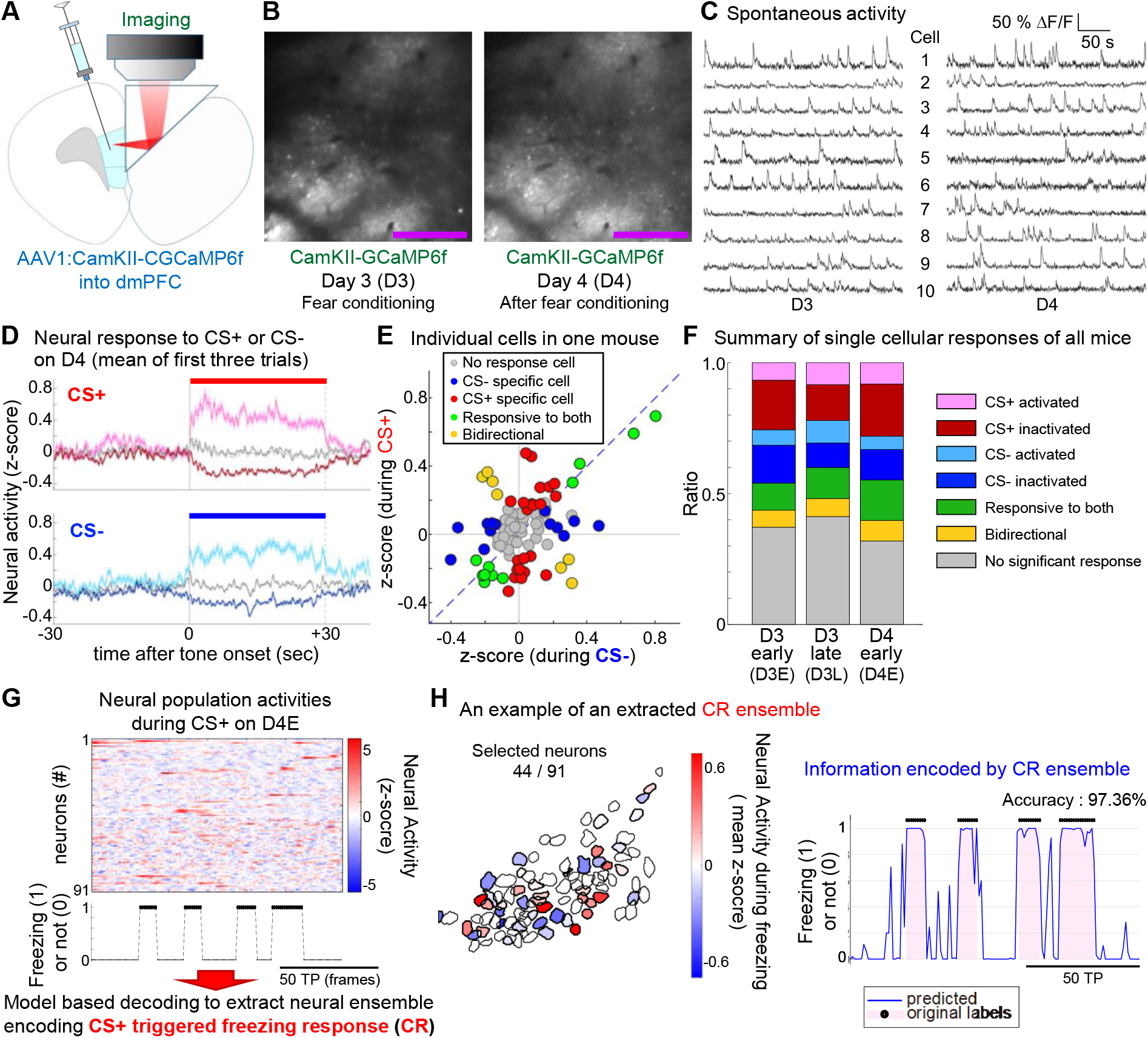
Longitudinal in vivo imaging in dmPFC and extraction of the neural ensemble encoding conditioned responses. (**A**) Microprism implantation along the midline for optical access to the dmPFC without cutting nerves. To visualize activity of excitatory neurons in the dmPFC, GCaMP6f was expressed by the AAV under the regulation of the CamKII promoter. (**B**) In vivo two-photon microscopy to detect single-cell neural activity visualized by GCaMP6f, chronically (day [D] 3 and D4) from the same set of neurons observed through the prism. See also Movie S1 and 2. Scale bar, 250 μm. (**C**) Traces of spontaneous Ca^2+^ activity from 10 example neurons in dmPFC, chronically on D3 and D4. (**D**) Summary of neural responses during the retrieval session (D4-early [D4E], mean of first three trials) to the CS+ or CS−. Mean of neural responses in each category (significantly activated [bright red or blue], inactivated [dark red or blue], and others [dark gray]), as well as the mean of all cells (light gray) are plotted. (**E**) Scatter plot showing responses of individual neurons to the CS+ and CS− in an example mouse during D4E. Each dot represents the mean response of each neuron. Blue, red, and green colors indicate that cells had a significant response as described in the panel. These features for all the mice are summarized in panel E. (**F**) Summary of response profiles at each phase (D3E, D3-late [D3L], and D4E, respectively; N=7 chronically recorded mice). (**G**) Schematic diagram showing how we extracted the CR ensembles. See the Materials and Methods for details. (**H**) An example of the CR ensemble and encoded neural representation of the behavior. (left) Extracted neurons are drawn with a bold margin, and the mean activity during CR (freezing) is shown in color. (right) Time course changes of neural representation encoded by the CR ensemble shown in the left panel. Black dots on the top of the graph and pink color in the graph indicate the timing of the actual CR, while the blue line shows information decoded by this CR ensemble. The plots show a part of the whole length of the data, and overall decoding accuracy was 97.36% in this example. TP, time points (i.e., image frames).

Prior to investigating population coding in the dmPFC, we first summarized the single-neuron responses to the CS+ and CS− before and after acquisition of the fear memory (Figs. 2D-F and S1). We found that approximately 60% of neurons exhibited a significant change in neural activity during the CS+ and/or CS−, and approximately 20% of neurons showed significant responses to both the CS+ and CS−. The distributions of these types of neurons were consistent throughout the learning process (Figs. 2F and S1). This type of “mixed selectivity” (responsive to variable task-relevant aspects) has been reported in the primate PFC (*23*) as well as in the mouse caudal mPFC during a decision-making task (*24*). The potential advantage of the mixed selectivity was proposed to enhance the number of tasks that each neural circuit, with a limited number of neurons, can handle, through high-dimensional neural representations implemented by a population of neurons (*23*, *25*). This encouraged us to further analyze the population coding for fear memory.

### Newly emerged and unique neuronal ensembles in dmPFC encoding conditioned responses

Our goal in this study was to dissect the computational architecture composed by a neural population in the dmPFC enabling the distinctive acquisition of a novel associative memory. For this purpose, we first extracted a group of neurons encoding the conditioned response (named CR ensemble) (Fig. 2G, H), and compared it with the neurons encoding regular locomotion (RL) to evaluate the uniqueness of the CR ensemble (Figs. 3 and S2). As methods to analyze neural architecture embedded in the neural population activities, previous studies utilized unsupervised algorithms such as Principal Component Analysis or Non-Negative Matrix Factorization (*14*, *26*, *27*). These algorithms first seek and distinguish embedded structures in neural data without considering the behavioral labels (e.g., freezing responses), which are further used to test which extracted structure is most likely to correlate with and explain respective behaviors, e.g., behavior A and B. By such methods, the identified ensembles corresponding to behavior A and the identified ensemble corresponding to behavior B may become dissimilar from each other as a result of methodological bias, irrespective of the actual neural architectures. In the present study, instead of these methods, we introduced a supervised and model-based decoding algorithm, named elastic net (*28*) (Figs. 2G-H). The elastic net is a regularization and variable selection algorithm based on the regression model (Fig. 2G; see the Materials and Methods for details) (*28*). This method enabled us to independently extract the CR ensemble and RL ensemble from the same mice (Figs. S2), and is thus helpful for further comparing the ensembles and systematically verifying whether neurons in the CR ensembles were unique or mostly overlapped with the RL ensembles.

**Fig. 3.**
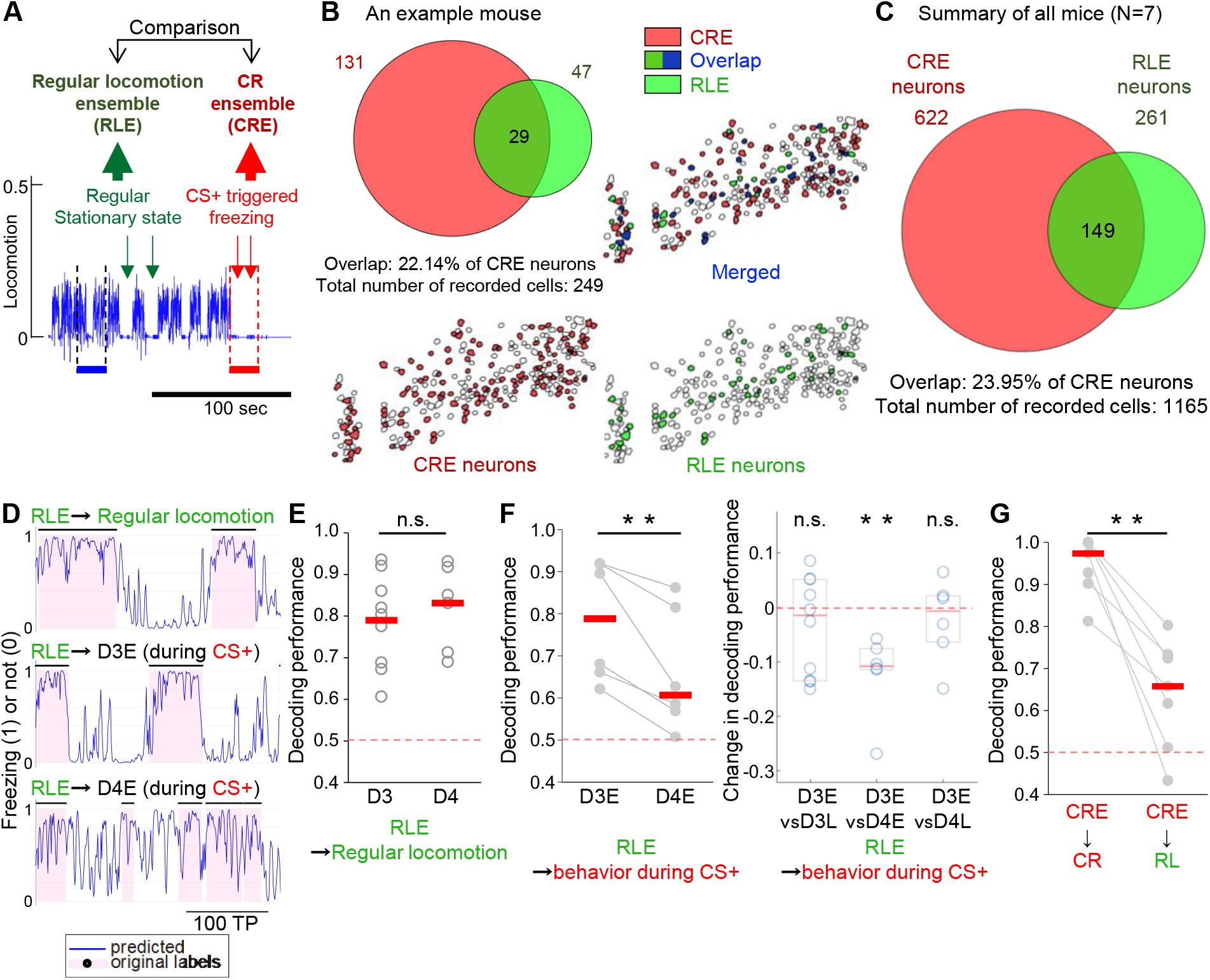
Emergence of unique CR ensembles after fear conditioning. (**A**) The CR ensemble was compared with the RL ensemble to evaluate the uniqueness. (**B**) An example Venn diagram and an example spatial map showing the overlap between CR ensemble neurons and RL ensemble neurons in an example mouse. (**C**) Summary of the overlap between the CR ensemble neurons and RL ensemble neurons of all mice (N=7, n=1165 neurons). (**D-F**) Decoding locomotion during regular locomotion (inter-trial interval) or during CS+ by the RL ensemble. (**D**) In an example mouse, an RL ensemble (RLE) that showed high accuracy for decoding performance to predict RL (top) also showed high decoding performance in predicting locomotion during CS+ at day 3-early (D3E). But the performance dropped when locomotion during CS+ at D4E (i.e., during fear retrieval) was also predicted by the RL ensemble. (**E**) Original decoding performance of the RL ensembles (i.e., predictability for RL) were not significantly different between D3 and D4. (**F**) (left) Decoding performance of RL ensembles to locomotion during CS+ at D4E (i.e., during fear retrieval) was significantly lower than that for D3E (i.e., before memory consolidation). (right) The change in decoding performance was systematically evaluated. Decoding performance was not significantly different between D3E and D3-late (D3L), or between D3E and D4L. (**G**) Decoding locomotion during CS+ by CR ensembles. Decoding performance was significantly decreased when the CR ensembles were applied to predict RL. Within D3, N=10; D3 vs D4 and within D4, N=7 pairs. A non-paired comparison (Wilcoxon rank sum test) was performed for panel D, while for the other comparisons in E and F, a paired permutation test was performed. For the decoding performance, we plotted the accuracy scores, while the AUC was very similar as shown in Fig. S5. **p<0.01; n.s., not significant. Red bars, median; box in panel E (left) indicates 25th and 75th percentiles.

We extracted CR ensembles using neural activity data obtained during the CS+ presentation of D4E (retrieval session), and evaluated the fitting and decoding performance of the obtained model (Fig. 2H) after optimizing the hyper parameter “alpha” for the elastic net as explained below (see the Materials and Methods for details). Compared with the conventional regularization algorithm Lasso, the advantage of the elastic net is that this procedure enables us to optimize the size of the selected population, especially when an analyzed neural network includes strongly correlated neural pairs, which is likely the case for our data considering the results shown below and previous electrophysiological observations (*14*). To carefully verify the overlap between the CR ensemble and RL ensemble, it is important to avoid missing neurons encoding the respective information. For this purpose, we evaluated remaining information encoded by neurons excluded from the CR ensemble at each alpha, and defined optimal alpha as the one that minimize such remaining information (Figs. S2C, D and S3; see also Materials and Methods). A wide range of the alpha values for the CR ensemble of each individual mouse was tested (Fig. S3). This systematic optimization procedure revealed a general trend that a larger alpha tended to select a smaller number of CR ensemble neurons (Fig. S3B, top), and though the decoding performance of the smaller number of selected CR ensembles was very high, equivalent to that of the others (Fig. S3B, middle), the removal of such a smaller portion from the whole set of neurons was not sufficient to substantially diminish the information encoded by the remaining neurons for some mice (Figs. S3B, top and bottom, and S3D-F). This suggests that the CR was redundantly encoded in the dmPFC. On the other hand, alpha values didn’t clearly affect the discrimination of the RL ensemble (Fig. S4). After determining the optimal alphas for individual circuits, we confirmed a substantial reduction of the decodability by removing all the selected CR ensemble neurons (Figs. S2D, and S3) and verified that a sufficiently large portion of the dmPFC neurons encoding the CR were selected as the CR ensemble neurons.

We eventually confirmed that the CR ensemble obtained by the optimal alpha was highly predictive for the CR during the retrieval session (see the example shown in Fig. 2H; mean ± SE of the prediction accuracy, 0.9450 ± 0.0265, N=7 mice; individual data are shown later in Fig. 3G). As for the spatial distribution, the identified CR ensemble neurons were spatially intermingled over the field of view, as shown in Figs. 2H and 3B.

Following the optimization of the hyper parameter alpha, we evaluated the specificity and uniqueness of the extracted CR ensemble. We confirmed that most of the neurons involved in the CR ensemble were unique and did not overlap with the RL ensemble (Figs. 3A-C). We then conceived the hypothesis that the unique CR ensemble might dominantly and exclusively explain the behaviors of the mice during CS +-evoked memory retrieval as an encoder of the acquired associative memory. If this is true, RL ensembles, distinct from CR ensembles (Fig. 3C), should have diminished decodability for the behavior during CS+ during memory retrieval. To test this possibility, we checked the decoding performance of the RL ensembles for the behaviors observed during the CS+ at each of the learning steps (Figs. 3D-G and S5). The decoding performance by the RL ensemble to the RL was similar between pre- and post-memory consolidation (Fig. 3E). The decoding performance of the RL ensemble to the behaviors during CS+ presentation at D3E (before fear memory consolidation) was similar to that for the RL (Figs. 3D, F). In contrast, the decoding performance of the RL ensemble to the behaviors during the CS+ on D4E (during fear memory retrieval) was significantly reduced compared with that of D3E (Figs. 3D, F). There was a small, but not significant, change during the fear conditioning (D3E vs D3L; Fig. S5), and importantly, the reduced decodability of the behavior during CS+ at D4E (memory retrieval) was substantially recovered after the extinction training (no significant difference between D3E and D4L, and a significant difference between D4E and D4L; Figs. 3F and S5). On the other hand, the decodability of CR ensembles was specific to the CR and not applicable to the RL on D4 (Fig. 3G). These results established that the CR, or the behavior during the memory retrieval, was dominantly explained by the CR ensembles, supporting the idea that the CR ensemble systematically extracted was a dominant and specific group of neurons encoding the CR during memory retrieval, emerged after consolidation of the fear memory and was suppressed by extinction.

### Coactivity within the CR ensemble was specifically enhanced after fear conditioning

In these CR ensembles, we observed a slight but significant increase in CS+ activatable neurons, but no change in CS+ inactivated neurons after fear conditioning (Fig. S6). In contrast, other cells (neurons that were not included in the CR ensembles: Non-CR ensemble [Non-CRE] neurons) exhibited no significant changes in the CS+ activatable neurons, with a significant increase in CS+ inactivated neurons. Neurons in the RL ensembles did not exhibit any significant change in CS+ responsiveness. We detected no significant change in CS− responsiveness in any of the categories. Because these CR ensembles were discriminated by the data and behavioral labels during the CS+, not by comparisons between those during the CS+ and those during the presentation of other stimuli, our method produced no bias toward the CS+ during selection of the CR ensemble neurons. These results indicated that there might be some mechanism that makes neurons involved in the CR ensembles dominantly activated by the CS+ after memory consolidation.

To further analyze and characterize the identified CR ensemble toward elucidating the mechanism underlying associative learning, we measured the change in the coactivity of the neural network during the CS+ presentation by comparing the pairwise correlation coefficients (R) (*29*) between pre- and post-memory consolidation. We found that, after the fear conditioning, only the positively correlated fraction was enhanced specifically within the CR ensemble, and not in the outside network (Non-CRE) (Fig. S7A). Statistical analyses demonstrated that this enhancement in positive correlation after the fear conditioning, as well as the enhanced ratio of significantly and positively correlated pairs, specifically occurred in the CR ensemble (Figs. S7A-C). Analyses based on the shuffled data, where the activity of each neuron was preserved but the temporal order was randomly shuffled neuron by neuron, revealed no significant difference between the CR ensemble and Non-CRE (Figs. S7A, C), suggesting that the specific enhancement of the coactivity of the CR ensemble in the real data did not derive from the enhanced neural activation. Similar enhancement of the coactivity was observed in the CR ensemble excluding the RL-ensemble overlapped neurons (Figs. S7A-C). In addition, changes in the coactivity across the categories (coactivity between CR ensembles and Non-CRE) were significantly smaller than those within the CR ensembles (Fig. S7C). These results led us to hypothesize that the functional connectivity within the CR ensemble was specifically enhanced as a result of the fear conditioning, contributing to enhance the coactivity.

### Enhanced internal connectivity and association with conditioned stimuli (CS) in the CR ensemble after fear conditioning

To test the hypothesis above, we introduced a probabilistic graphical model method, the conditional random field (CRF) model (*30*, *31*). This method evaluates the conditional probability that a group of neurons fire together given that one neuron is active (Fig. 4A). Among the various mathematical algorithms used to evaluate possible functional connectivity of neural networks and ensembles, the CRF model is one of the most reliable methods because the results of the calculation (functional connectivity) have already been carefully evaluated by two-photon holographic optogenetics and consequential behavioral modulation (*30*, *31*).

**Fig. 4.**
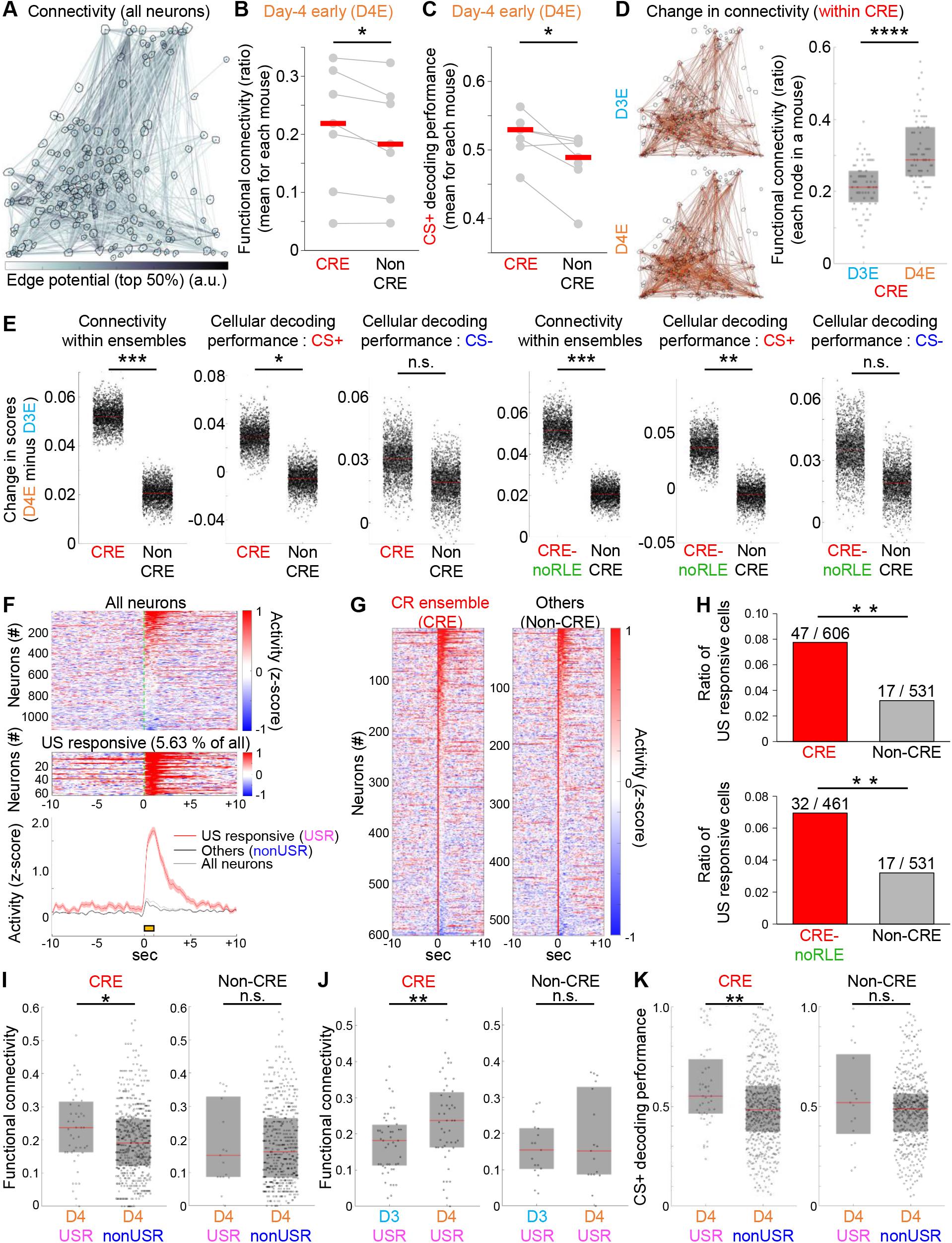
Enhanced functional connectivity and CS+ predictability in the CR ensemble with an emergent hub of US responsive neurons after fear conditioning. (**A**) Functional connectivity between neurons in an example circuit. Among all the possible connections for all pairs of neurons, the CRF model enables the estimation of functional connections, as well as the dependencies of connected pairs. In this panel, the top 50% edge potentials were visualized. (**B**) During day 4-early [D4E], the functional connectivity within CRE was significantly higher than that of Non-CRE. (**C**) During D4E, the predictability for CS+ in CRE was also significantly higher than that in Non-CRE. (**D**) Change in functional connectivity within CRE of an example circuit. This is the same as the circuit shown in A, but only the connectivity of the CRE neurons marked by the red ellipses were analyzed. Left panel shows the change in the connectivity between day 3-early [D3E] and D4E, while the right panel shows the change in the ratio of functional connectivity per all possible connections for individual nodes (i.e., individual neurons). (**E**) Summary of changes in functional connectivity and cellular decoding performance for CS+ and CS− of all observed networks (N=7 mice). Differences (D4E minus D3E) of these scores are plotted as a result of bootstrap resampling (2000 times) to compare CRE and Non-CRE, or CRE-noRLE (CRE neurons excluding those overlapping with RL ensemble neurons) and Non-CRE. (**F**) A part of the recorded neurons in the dmPFC showed increased activity upon US presentation on day 3 (D3) during fear conditioning. Mean activity over 7 trials of all (top) or US-responsive (middle) neurons, and the mean ± s.e.m. of respective categories (bottom) are plotted. Green dotted line indicates the onset of the US, and yellow bar indicates the 1-s duration of the US presentation. (**G**) Summary of US responses of CR ensemble neurons (CRE) and others (Non-CRE). All individual neurons for the respective categories are plotted. (**H**) Neurons responding to the US on D3 were predominantly involved in the CRE on D4 after the fear conditioning. The difference between CRE vs Non-CRE, as well as CRE-noRLE vs Non-CRE, was statistically evaluated. (**I**) Comparison of functional connectivity between US responsive neurons (USR) and others (nonUSR) on D4. In the CRE network, USR became more connected within the network than nonUSR, while there was no significant difference between USR and nonUSR outside of the CRE (Non-CRE). (**J**) The higher connectivity of USR on D4 was experience-dependent. Functional connectivity of USR on D4 was significantly higher in CRE, while there was no significant difference between them in Non-CRE. (**K**) USR in the CRE exhibited significantly higher decoding performance of CS+ than nonUSR, which was not the case in Non-CRE. A paired permutation test was used for the statistics in B and C. The Wilcoxon signed-rank test was used for the statistics in D. The data obtained by bootstrap resampling were statistically analyzed as described in the Materials and Methods. Because the number of USR was limited (only 5.63% under the present definition), the analyses shown in panels F-K were performed with data pooled together from all mice (N=7 mice). Fisher’s exact test was used for the statistics in H, a non-paired comparison (Wilcoxon rank sum test) was used in I and K, and the Wilcoxon signed-rank test was used in J. *p<0.05; **p<0.01; ***p<0.001; ****p<0.0001; n.s., not significant. Red bars, median; gray boxes in panels D, E, I-K indicate 25th and 75th percentiles.

Using this method, we found that, after the fear conditioning (D4E), the functional connectivity was significantly higher in the CR ensemble (Fig. 4B). This method also allowed us to evaluate the information coding of any arbitrary label, e.g., CS+ or CS−, and we found that the CS+ information encoded by the CR ensemble was significantly higher than that of Non-CRE (Fig. 4C). Importantly, our method did not produce any bias to the CS+ in selecting CR ensemble neurons, as explained above. Therefore, this result indicates that the neural population encoding the CR was dominantly associated with the CS+ information. In addition, we found that the enhancement in both the functional connectivity and CS+ predictability was experience-dependent and derived after the fear conditioning, dominantly in the CR ensemble neurons (Figs. 4D, E). In contrast, the changes in information coding for the CS− were not significantly different between the CR ensemble and the Non-CRE (Fig. 4E). Therefore, the emergence of the CR ensemble after fear conditioning was accompanied by the enhancement of the internal coactivity, functional connectivity and association with the CS+ selectively within the CR ensemble neurons, indicating that fear conditioning drives dmPFC reorganization to generate novel associative circuits for CS− to-CR transformation.

### Neurons responding to US during fear conditioning anterogradely became a hub of the CR ensemble

Finally, we hypothesized that the dmPFC reorganization that we observed after fear conditioning might occur via activity-dependent modulation during the repeated CS+-US pairing. This led us to search for the signature of this plasticity.

During the fear conditioning, we observed that some of the dmPFC neurons strongly responded to the US (Fig. 4F). Interestingly, statistical analyses demonstrated that neurons responsive to the US during fear conditioning were predominantly and significantly more involved in the CR ensemble after the fear conditioning (Figs. 4G, H). This suggests that these US-responsive neurons (USR) were preferably integrated into the CR ensemble, in which functional connectivity might also be modulated and strengthened by US-evoked activity, perhaps together with the paired CS+ signal.

To test this possibility, we performed further analyses based on the CRF modeling. We found that the USR became functionally more connected within the CR ensemble than non-US responsive neurons, while these differences were not observed in Non-CRE (Fig. 4I). This higher connectivity was a result of the fear conditioning (Fig. 4J). The information coding for the CS+ was also significantly higher in the USR, specifically in the CR ensemble (Fig. 4K), suggesting that the US-responsive network was dominantly associated with the CS+ network when it became integrated into the newly emerged CR ensemble. According to a previous study, higher functional connectivity and higher decoding performance of sensory stimuli are typical features of pattern completion cells whose activation could efficiently enhance the entire ensemble activity for a specific sensory stimulus and promote the stimulus-associated behaviors of mice (*30*). These results collectively suggest that the USR in the dmPFC became a hub of the novel neural ensemble linking the CS+ to the CR, a memory-evoked behavior, after the repeated CS+ and US parings.

## Discussion

Altogether, our results based on the combination of methods for computational dissection and longitudinal recording in the dmPFC demonstrate learning-dependent dynamic modulation of population coding for associative fear learning, structured on an activity-dependent hub-network formation within the dmPFC. Through regularized regression methods and graphical modeling, we found that the repeated CS+-US pairing for the associative learning drives the dmPFC reorganization to generate novel and unique neural circuits for CS-to-CR transformation, with enhanced internal coactivity, connectivity, and association with the CS+. Upon this prefrontal reorganization, neurons activated by the US during fear conditioning were anterogradely and predominantly integrated into the CR ensemble. The eventual network stemming from these USR gained typical features of pattern completion cells of the CR ensemble, which are supposed to work as a hub in the prefrontal networks to predominantly relay the CS+ information and promote the CR (Fig. S8).

To our knowledge, this is the first in vivo evidence directly demonstrating that the prefrontal neural circuit for the associative memory was actually built based on an enhanced association between the US network and the CS+ network as a result of CS+-US pairing and triggered network reorganization. More than 60 years ago, Hebb proposed that repeated coactivation of a group of neurons might create a memory trace through the enhancement of connections (*32*). Our results suggest that Hebbian plasticity (i.e., fire together, wire together) might underlie the reorganization of the prefrontal network structure during associative learning, enabling the emergence of a strong link between the US signaling pathway and the CS+ signaling pathway to form a novel CR circuit.

CR information was redundantly encoded in the dmPFC. The advantage of the redundancy is not clear, but because fear memory is critical for animal survival, it is possible that the redundant coding for the fear memory is not inefficient but rather evolutionarily crucial. On the other hand, although the redundancy can also be considered inefficient in terms of the short-term cost, because the dmPFC is known to be involved in long-term memory via brain-wide networks (*12*, *33*, *34*), it would be interesting to investigate whether the redundantly encoded information for the CR is maintained or diminishes by longer-term continuous recording, and whether it is related to the brain-wide regulation of memory using virus-based anterograde or retrograde fluorescent labeling techniques to simultaneously dissect the downstream or upstream structures.

As we have successfully discriminated the specific neural population encoding the CR as well as the detailed internal structure with a hub of the US-responsive neurons, further testing the causality of the identified structure to behavior by holographic optogenetics (*30*) could be intriguing. But importantly, we also found that the dmPFC responds to auditory signals even before memory consolidation (Figs. 2D-F, S1) and that the CR ensemble predominantly includes the US-responsive neurons (Fig. 4F). Because enhancing the sensory coding can modify behavioral responses in a task based on the sensory stimuli as demonstrated before (*30*), and because activating US-responsive neurons may sufficiently encourage defensive freezing behaviors as unconditioned responses, further mathematical dissection and additional anatomical dissection discussed in the preceding paragraph would be the next important step to more precisely identify the memory-specific connections and information flow to be tested by the holographic optogenetics.

## Acknowledgments

We thank M. Sato, Y. Hayashi, M. Matsuzaki, Y. Masamizu, K. Takao, K. Miyamichi, A. Yamanaka, K. Seiriki, J. Chikazoe, H. Tsukada, D. Peterka, A. Packer, A. Noritake, D. Cheung, and T. Imai for technical advice and discussion. We thank the Bionanophotonics Consortium under the MEXT Scientific Research on Innovative Area “Spying Minority in Biological Phenomena” (Grant Number, 23115001) for assistance in microscopy, and the ISIR Machine shop for development of the running disk system. We also thank the members of the laboratory for their help, especially T. Matsuda and T. Wazawa for technical advice, and T. Kobayashi for help with the mice.

## Author contributions

M.A. conceived and coordinated the whole project. M.A. designed and performed behavioral experiments with the support of A.K., H.H., and T.N.; M.A. constructed in vivo imaging system and performed imaging experiments with the support of Y.A. and T.N.; M.A. performed data analyses with the support of I.S, Y.R.T., L.C.-R., M.Y., T.I., and J.N.; M.A. wrote the paper, with contributions from all authors.

## Data and materials availability

The data that support the findings of this study are available from the corresponding author upon reasonable request. Custom codes used to analyze data in this study are available from the corresponding author upon reasonable request.

## Materials and Methods

### Animals

All animal experiments were carried out in accordance with the Institutional Guidance on Animal Experimentation and with permission from the Animal Experiment Committee of Osaka University (authorization number: 3348), or in accordance with National Institutes of Health guidelines and approved by the National Institute for Physiological Sciences Animal Care and Use Committee (approval number 18A102). Male C57BL/6 or PV-Cre mice (Jax: 008069) mice housed under a 12-h light/dark cycle with free access to food and water were used for all experiments. Behavioral experiments were performed during the dark cycle (i.e., when mice were normally awake) using single-housed mice. Mice at 4–6 months of age were used for the behavioral and imaging experiments.

### Virus injection

To express GCaMP6f, a genetically encoded calcium indicator to monitor the neural activity, we used a gene expression system based on the AAV vector. Viruses were injected into mice at postnatal day (P) 50-120 for in vivo experiments, at least 1 month before the microprism implantation, which was followed by the in vivo experiments 1–3 months after the implantation. Injection procedures were performed as described previously (*29*), with some modifications. During surgery, the mice were anesthetized with isoflurane (initially 2% [partial pressure in air] and then reduced to 1%). A small circle (~1 mm in diameter) of the skull was thinned over the left mPFC using a dental drill to mark the site for a small craniotomy. AAV1/CamKII.GCaMP6f was obtained from the University of Pennsylvania Vector Core, and injected into the left mPFC (slightly away from the imaging target area to avoid damaging the field of view) at three sites (depth 1.0, 1.5, and 2.0 mm from the pial surface, volume 375 nl/site) to cover the dorsal mPFC, over a 5-min period at each depth using a UMP3 microsyringe pump (World Precision Instruments). The X-Y coordinates for the injection site was usually 0.5 mm lateral to the midline and 2.0 mm rostral to bregma, but if large blood vessels obstructed the position, we shifted the insertion site slightly to avoid the vessels. The beveled side of the injection needle was faced to the midline so that the needle could be smoothly inserted and the virus would cover the surface layers of the mPFC. We designed our injection protocol (especially the volume and depth) carefully to widely cover the mPFC areas, while the anatomical coordinates of the field of view for the two-photon imaging were precisely targeted using the position of the pial surface and the sinus, which were usually visible through the imaging window prepared as shown below, as a guide (the field of view ranged from a depth of ~0.9-1.9 mm and centered at a depth of ~1.1-1.5 mm from the pial surface and the sinus).

### In vivo two-photon imaging

In vivo two-photon imaging was performed as described previously (*19*, *29*), with modifications to pair with our new experimental system. At 1–3 months after the virus injection, the mice were anesthetized with isoflurane (initially 2% [partial pressure in air] and reduced to 1%). A titanium head plate described in a previous paper by Goldy et al. (*35*) was selected for the present study to minimize the area laying over the ear and to minimize the blockage of auditory input through the ear. The head plate was attached to the skull with dental cement. For the subsequent microprism implantation, a square cranial window (~2.3 x 2.3 mm) was carefully made with minimal bleeding above the right mPFC, the hemisphere opposite to the virus injection site. An implantable microprism assembly(*19*), comprising a 2-mm right angle glass microprism (TS N-BK7, 2mm AL+MgF2, Edmund) bonded to a 2×2 mm square cover glass (No.1; Matsunami) for the middle position and a 4×4 or 3×4 mm glass window at the surface position of the imaging window, was prepared and inserted into the subdural space within the fissure along the midline as described previously(*19*) to avoid harming any nerves surrounding the mPFC network in both hemispheres, allowing for visualization of the left mPFC, which was previously injected with the GCaMP6f virus, through the imaging window. The area directly beneath the microprism was compressed but remained intact. This insertion procedure sometimes caused a small amount of bleeding that covered the imaging site, but even in that case, the imaging window became clear after waiting at least a month before performing the experiments. As reported before (*19*), the mice recovered quickly and displayed no gross impairments or behavioral differences compared with non-implanted mice, enabling chronic imaging of the dmPFC in behaving mice.

The activity of dorsal mPFC neurons was recorded by imaging fluorescence changes with a FVMPE-RS two-photon microscope (Olympus) and a Mai Tai DeepSee Ti:sapphire laser (Spectra-Physics) at 920 nm, through a 4x dry objective, 0.28 N.A. (Olympus) or a 16x water immersion objective, 0.80 N.A. (Nikon). Mean (±SE) frame rate was 8.96 ± 0.87 (frames/s). GCaMP6f signals were detected via the band-pass emission filter (495-540nm). As the GCaMP6f was expressed under the regulation of the CaMKII promoter (*21*, *22*), all of the recording targets were assumed to be excitatory neurons (*20*). Scanning and image acquisition were controlled by FV30S-SW image acquisition and processing software (Olympus). To smoothly set the mice below the objective lens for the imaging, light and minimal-duration isoflurane (2.0% for less than 2-3 min) anesthesia was used, and behavioral and imaging experiments were started 5 min after the mice awoke and began locomoting on the running disk, which was visually confirmed via the video camera (VLG-02, Baumer) under infrared light-emitting diode illumination (850nm: LDL-130X15IR2-850, CCS Inc.). To detect neural activity from the same set of neurons in each mouse over multiple days, the depth from the surface of the brain (dmPFC area) and configuration of blood vessels and basal GCaMP6f signals in each field of view were recorded and referenced as described previously (*36*).

### Fear conditioning, memory retrieval, and extinction under the microscope

The experiments were designed according to previous studies, with some modification to optimize conditions for the two-photon microscope system (*13*, *14*, *18*). The heads of the mice were fixed under the objective lens for two-photon imaging, allowing them to run freely on the running disk placed below them, and locomotion and the freezing response were measured by the rotation of the running disk, as previously described (*37*). Experiments were performed in a completely dark environment to protect the detector (photo multiplier tube) for the two-photon imaging from the room light. We prepared two different types of running disks to establish two different contexts, as used in conventional fear conditioning experiments for head-unfixed mice (*13*, *14*, *18*). Disk A was made of light-colored plastic with ridges from the center to the rim that the mice could grip to allow them to easily rotate (and walk on) the disk (*37*). Disk A was used for adaptation (D1 and D2) and for retrieval and extinction (D4). Disk B was built for the fear conditioning (D3), and comprised a grid made of stainless steel bars (Fig. 1A), which was attached to a foot shock generator (SGA-2010, O’HARA & CO., LTD) via an electrical slip ring so that electrical current to this running disk for the foot shock (US) could be stably delivered to the mouse irrespective of whether the running disk was rotating. The behavioral sessions on each day began only after the mouse was constantly locomoting for more than 5 min. The running disks and the surrounding area (inside the cage for the microscope) were cleaned with 70% ethanol before and after each experiment. To score freezing behavior, the speed of the mouse locomotion was measured by the rotation speed of the running disk (*37*), and mice were considered to be stationary (during no CS presentation) or freezing (during CS+/retrieval) if no movement was detected for at least 1 s. On D1 and D2, the mice underwent an adaptation session with disk A for an hour each day, to familiarize them with the novel environment. On D3, the mice underwent a habituation session in context B, in which they received four presentations of the CS− and CS+ alternately (total CS duration, 30 s for each trial; consisting of 50-ms pips at 1 Hz repeated 30 times; pip frequency, 7.5 kHz or white-noise, respectively, 80-dB sound pressure level (60-dB basal room noise produced by the air conditioning system, and 20-dB for the CS)). The habituation session was immediately followed by discriminative fear conditioning (*13*, *14*, *18*) on the same day by pairing the CS+ with a US (1-s foot shock, 7 CS+−US pairings).The intensity of the foot shock was usually 0.05~0.1 mA, but when mice showed no responses at all, which was probably caused by that a part of the running disk became dirty or wet by mice and the foot shock might be suppressed by this during the experiment, an intensity of 0.25~0.45 mA was used. The onset of the US coincided with the onset of the last sound pip of each 30-s CS trial. The CS−and the CS+ trials were performed alternately (inter-trial intervals, 50–150 s). On D4, conditioned mice underwent a retrieval session followed by an extinction session on disk A during which they received 4 presentations of the CS− and 12 presentations of the CS+. During the experiment (D1-4), the mouse was continuously encouraged to locomote by administering a 4-ul drop of saccharin water per 100 cm of locomoting, provided through a spout placed near their mouth (*36*) so that the freezing response could be discriminably detected as decreased locomotion (Fig. 1). The mice were not water-deprived. The locomotion speed and timings of the tones and the foot shock were synchronously recorded with image acquisition (GCaMP6f imaging in dmPFC) using NI software (Labview; National Instruments) and NI-DAQ (National Instruments). The results shown in Fig. 1 show that this protocol led to the mice successfully learning the CS+-US association, and show a reduction in locomotion in response to the CS+, but not the CS−, and not before but only after the fear conditioning session, enabling us to observe changes in neural representations in the dmPFC as a result of the fear conditioning.

### Imaging data analyses and statistics

The raw images of the GCaMP6f signals in the dmPFC were processed to correct for brain motion artifacts using the enhanced correlation coefficient image alignment algorithm (*38*). To apply the same regions of interest (ROIs) for analyzing the images obtained across multiple days, the movies from the same mouse were precisely aligned with each other using the same enhanced correlation coefficient algorithm as above, while, for a local shift (shift of a few pixels in a small number of neurons among all recorded cells), the corresponding ROIs were manually adjusted.

The ROIs for the detection of neural activity were automatically selected using a constrained nonnegative matrix factorization algorithm in MATLAB as described previously (*39*), with some manual adjustment. Further steps to process the GCaMP6f signals for measurements of the signal change (ΔF/F) of each neuron were performed as described previously (*29*, *40*); although the same constrained nonnegative matrix factorization package for ROI detection also provides an option for signal processing that was not sufficiently optimized to analyze our data, which were obtained over several days with more than 30,000 frames each day. Fluctuations in the background fluorescence, which contains synchronous fractions across nearby neurons (*39*, *40*), was subtracted before calculating the ΔF/F of GCaMP6f signals as described previously (*29*). Briefly, a ring-shaped “background ROI” was created for each ROI 2–5 pixels away from the border of each neuronal ROI to a width of 30–35 pixels, and the size was adjusted to contain at least 20 pixels in each background ROI after completing the following steps. From the background ROI, we removed the pixels that belonged to any neuronal ROIs, and the ROIs that contained artificially added pixels (black pixels added at the edge of the image due to the motion correction procedure) at any time-point. We then removed the pixels that, at some time-point(s), showed signals exceeding that of the neuronal ROI by two standard deviations of the difference between each background ROI pixel time series and the neuronal ROI time series. The resulting background ROI signals were averaged at each time-point, and a moving average of the time series was calculated. Using the moving average instead of the raw background ROI signal was helpful to minimize the production of an artificially large increase or decrease at each time-point due to the subtraction, which could have altered the analyses of the timing of neural activations. Pixels within each neuronal ROI were also averaged to give a single time course, and then the background ROI signal was subtracted. Then, the ΔF/F of GCaMP6f signals of all neurons in each circuit was calculated. For most of the analyses and comparisons of the results from multiple mice, the ΔF/F data were further z-normalized within each experiment (same mouse, same day) as described previously (*13*, *18*). On the other hand, particularly for the CRF modeling used to evaluate the functional network connectivity, the spike probabilities were inferred from the ΔF/F as an alternative estimate of neuronal activation using a constrained sparse nonnegative calcium deconvolution method (*39*). We used the code “constrained_foopsi.m” (*39*), and the parameters used in the calculation were not manually selected but estimated from the data by the code. After inference of the spike probability and further thresholding by two standard deviations, the obtained binominal data were further binned (bin size: 1 s). Importantly, the results obtained by CRF modeling were consistent with the results of the coactivity analyses based on the ΔF/F (and z-normalized ΔF/F) (Fig. S7), providing substantial support that the analyses based on both estimates complemented each other for the data analyzed in the present study. While neurons for the analyses were initially automatically detected, neurons responding to noisy signals with no apparent calcium transient at any time during the experimental days were identified by visual inspection and excluded from further analysis.

For the statistical analysis, we used MATLAB (MathWorks, Natick, MA). The Wilcoxon signed rank tests for paired comparisons or the Wilcoxon rank sum test (equivalent to Mann-Whitney U test) for unpaired comparisons was used to determine statistical significance (P < 0.05) unless otherwise indicated. Two-tailed tests were selected for all statistical analyses. All p-values less than 0.0001 are described as “P<0.0001” (or ****). Graphs were produced by MATLAB (MathWorks) or Excel (Microsoft). When comparing two groups (e.g. D3 vs D4) consisting of the results of multiple mice, in addition to the analyses using original data (e.g. N=7 vs N=7 [D3 vs D4]), we performed bootstrap resampling to more systematically estimate representative values (e.g. mean or median) of each mouse or each group where the number of recorded neurons in each field view varied. When statistically comparing original data (e.g. comparing D3 vs D4), we used a paired permutation test that does not require any assumptions regarding the data distribution, though the p-values obtained by this method and the evaluated statistical significance were very similar to those obtained by the paired t-test in almost all cases. For the analyses based on bootstrap resampling followed by statistical comparison, random resampling (with accepting overlapped sampling) from each mouse was performed in total with the same number as that of the original data of each mouse for each resampling round, and the means (e.g. of 7 mice each day) and the means of the difference or ratio (e.g. difference between D3 vs D4 averaged over mice) were calculated. This was repeated 2000 times to derive the distribution (of 2000 bootstrap replications) for each estimate, and the statistical significance was evaluated based on the 95% confidence interval.

In the present study, to compare changes in neural responses and ensemble representations before and after the fear memory consolidation without any bias, we did not exclude neurons that showed no response to the CS on D4 from the analyses, which was done in some previous experiments (e.g. (*18*)). Neurons for the analyses were automatically selected based on the neural responses, as described above, and all neurons that exhibited clear activity during at least one of the experimental days were included for the analyses irrespective of whether it was during the CS presentation or only during no CS presentation, considering the previous work suggesting that not only the neurons that typically respond to the CS, but also other types of neurons (including those of mixed selectivity) are important for population coding in the prefrontal network (*23*).

The significance of CS-induced neural responses was determined according to previous studies (*13*, *18*). Signals during CS presentation were normalized to baseline activity using a z-score transformation, as described previously (*13*, *18*). The CS-induced neural activity for each stimulus was then calculated as the mean of the activity during ~1 s from each stimulus onset (depending on the imaging frame rates, we set the number of frames to be used for this calculation so that sampling duration was closer to 1 s but the frames that overlapped with the next stimulus onset was excluded). The last sound pip of each 30-s CS trial was also excluded from this analysis because, during fear conditioning, the last sound pip of the CS+ overlapped with the US (we excluded the last pip data not only for analysis of CS+-evoked responses during fear conditioning but for all data analyses on both D3 and D4, for both CS+ and CS−). They were averaged over blocks of 3 CS trials consisting of 87 individual sound pips in total, for D3E (first three trials during the fear conditioning session), D3L (last three trials during fear conditioning on D3), D4E (first three trials on D4, as responses during fear memory retrieval), and D4L (last three trials only for CS+ on D4 as responses during extinction), respectively, or used to statistically test whether the responses of each neuron were significantly different from zero (baseline) and to define CS− activated / -inactivated neurons.

To define US responsive neurons, because the number of US were limited (7 stimuli in total for each mouse), the mean z-score of each neuron for 1.5 s from the US onset was calculated, and US responsive neurons were defined as neurons with responses of one standard deviation or larger. The number of USR was very limited (zero or only a few for some of the mice) as they were only around 5 % on average, and therefore all the analyses shown in Figs. 4I-K were performed with pooled data from all mice (N=7 mice).

To evaluate the coactivation of neural activity in the dmPFC network, we calculated cell-to-cell pair-wise correlations within each ensemble using Pearson’s correlation coefficient, from the GCaMP6f signals (z-normalized ΔF/F) of two cells over the duration of the CS+ presentation, as described before (*29*). The calculated correlation coefficients (R) were statistically analyzed. As a complementary analysis, we also used the inferred spike probability to analyze the functional connectivity, as explained in the section describing the CRF model, which revealed consistent results as shown in the results section. We further performed analyses based on surrogate datasets, as described in previous studies (*29*, *41*). For this, the total activity of each neuron was preserved, but only the timing was shuffled randomly within each neuron, followed by calculation of the correlation coefficients of shuffled data.

### Extraction of neuronal ensembles

To directly differentiate neural populations (ensembles) encoding the CR (i.e., suppressed locomotion triggered by CS+ during the memory retrieval) and those encoding RL (i.e., stationary or locomotive state during no CS presentation), we used the elastic net(*28*), a regularization and variable selection algorithm that enabled us to systematically extract neurons encoding respective target behaviors. For this, we used the “lassoglm” function of MATLAB R2019b. Because this method allowed us to identify different ensembles for different behaviors independently from the same mice, we used this to verify whether neurons in CR ensembles were unique or mostly overlapped with RL ensembles (Figs. 3 and S2). Compared with the conventional sparse modeling method called Lasso (least absolute shrinkage and selection operator), the advantage of the elastic net is that the hyper parameter “alpha” additively enables the adjustment of the size of selected neurons depending on the data; when the analyzed data include strongly correlated pairs, which appeared to be the case for our data as shown in Fig. S7, conventional Lasso removes redundant predictors and selects only one or a part of such a synchronous population, but in the elastic net, lowering the alpha value increases their inclusion, which is helpful toward preventing missing encoder neurons.

When extracting the CR ensemble, we used data only during the CS+ presentation of D4E (retrieval session) and identified neurons informative for distinguishing whether animals exhibited freezing behavior or were locomoting during the CS+ so that the auditory information of the CS was not considered for identifying the ensemble neurons. While mice exhibited the CR as suppressed locomotion during the fear memory retrieval session (Fig. 1), they also showed more or less locomotion intermittently, and both labels (freezing and locomotive) are required to perform the regression based on the elastic net; only the data containing at least 10% of each label (freezing and locomotive) were used to discriminate ensembles in the present study. On the other hand, for extracting the RL ensemble, we used data only during the no-CS presentation (for D3 and D4). Learning the elastic net is formulated as follows.

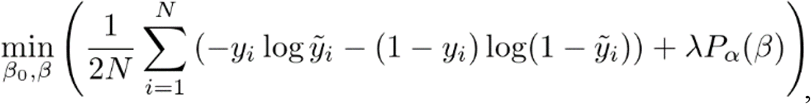

where

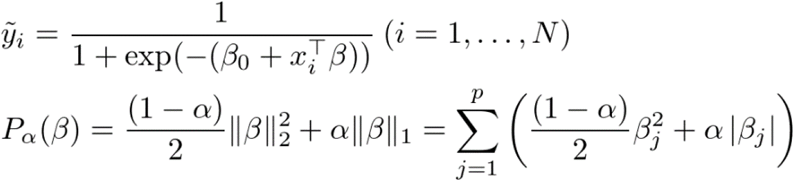

and N is the number of observations; y_i_ is the behavior (freezing/stationary y_i=1 or locomotive y_i =0) at observation i; x_i_ is data (neuronal activity), a vector of p values at observation i; λ is a positive regularization parameter; parameters β_0_ and β are a scalar variable and a p-dimensional vector, respectively. As λ increases, the number of nonzero components of β decreases. The elastic net is a hybrid of ridge regression and lasso regularization: when alpha (α) = 1, elastic net is the same as lasso, while, as α shrinks toward 0, elastic net approaches ridge regression. For other values of alpha (α), the penalty term Pα(β) interpolates between the L^1^ norm of β and the squared L^2^ norm of β. Lasso is sensitive to correlations between variables and can choose one if there are two highly correlated and useful variables, whereas elastic net is more likely to select both useful variables, which leads to more stable variable selection. The tuning parameter λ controls the overall strength of the penalty. β_j_ is the coefficient for the corresponding neuron j estimated by this model. Because this method is designed to sparsely leave the coefficients β_j_ for the respective neurons, we could identify neurons with a non-zero coefficient as ones of substantial decodability (i.e., ensemble neurons). The lambda value with minimum expected deviance, as calculated during cross-validation, was selectively used to define these beta coefficients for each dataset. To avoid an imbalance of the number of original labels for respective states (e.g. freezing or locomotive for CR ensembles) for the training, the same number of data points from respective states were randomly selected to prepare the training data despite an overlap, a total of 900 samples for each, and used to produce the model. We found that the eventual model and non-zero-coefficient neurons slightly varied trial by trial. To accurately define each ensemble, we repeatedly performed this procedure (random sampling and modeling) 100 times to obtain the distribution of each beta value. Gaussian fitting was performed to define the centroid and the 95% confidence interval of each distribution of each beta, and then the 95% confidence interval was used to determine whether or not they were significantly different from zero (enabling us to maintain sparsity), with the centroid being used to define the final beta values of non-zero coefficient neurons to build the model. To evaluate the fitting and decoding performance of the obtained model, the prediction accuracy and the area under the curve (AUC) of receiver operating characteristic curve (ROC) were calculated, respectively, revealing that those scores were very similar and highly correlated with each other (Fig. S5).

Based on the above-described procedure, we next optimized the alpha values. Ideally, if all the informative neurons can be extracted into the selected CR ensembles, the remaining neurons should have poor decoding performance. According to this idea, to optimize the alpha value, after building a model at each alpha for each mouse (“AUC original” in Fig. S3A), we compared the difference in decoding performance between “AUC CRE-rem” and “AUC nonCRE-rem” (Figs. S3A). AUC CRE-rem is the AUC value calculated by an elastic net model built with the neurons, excluding the original CR ensemble neurons. On the other hand, AUC nonCRE-rem is the AUC value calculated by the neurons, excluding neurons other than original CR ensemble neurons, randomly selected, and the number of excluded neurons was the same as the number of original CR ensemble neurons (so that the number of neurons used to calculate AUC nonCRE-rem were set to be the same as that used for AUC CRE-rem calculation). The “AUC difference” (Fig. S3A) between those two values was calculated to estimate the degree of remaining information, and in principle, we defined the best alpha based on the maximum AUC difference for each mouse independently. In addition, for further statistical evaluation to define the optimal alpha as explained below, we repeated these procedures 10 times for both “AUC CRE-rem” and “AUC nonCRE-rem”.

As shown in Fig. S3B, although the decoding performance of the original CR ensembles (i.e., AUC original in Fig. S3A) was not affected by the alpha (Fig. S3B, middle), the size of the CR ensemble was affected, and a smaller alpha generally resulted in a larger number of selected neurons for each CR ensemble (Fig. S3B, top), suggesting that the CR information might be redundantly encoded in the dmPFC as discussed in detail later. On the other hand, the influence of the alpha on the AUC difference was more complicated. As explained above, we defined the best alpha based on the maximum AUC difference for each mouse independently, but in some exceptional cases as shown in Fig. S3D (mouse #3), when the other alpha(s) showed a AUC difference(s) not significantly far from the maximum AUC difference, the alpha of the smallest of the ensembles among those alphas, i.e., largest alpha among them, was selected to avoid unnecessarily including additional neurons that did not improve the AUC difference (e.g. in mouse #3, alpha = 0.1, 0.05, 0.01 showed similar AUC differences and there was no statistically significant difference between them [Wilcoxon rank sum test, alpha of maximum AUC difference vs the other alpha, n=10 estimates for each calculated as explained above], so in this case, the largest alpha 0.1 among those three was selected to define the CR ensemble for this mouse).

These results revealed two important points. First, searching around the alpha value may be important in some cases. Considering this, we also searched alphas in the case of RL ensembles (Fig. S4), and found that there was no difference among the various alphas, for the RL ensembles, even if we tested an additional number of reference frames (means of the neural activities over the past or future several frames were used as neural activity data to predict a single label at each single time-point, which showed no significant difference from each other, evaluated by the Friedman test, a non-parametric statistical test similar to the parametric one-way repeated measures ANOVA). Therefore, in the present study, we fixed the alpha to define RL ensembles at 0.75 for most of the analyses, except for the data in Figs. S4 and S5, where we evaluated the influence of the alpha for RL ensembles.

Second, fear memory triggering the CR might be redundantly encoded in the dmPFC. As discussed above, although decoding performance of the original CR ensembles was not affected by the alpha (Fig. S3B, middle), the size of the CR ensemble was affected, and a smaller alpha generally resulted in a larger number of selected neurons for each CR ensemble (Fig. S3B, top). In addition, when the alpha was fixed at alpha (A) =0.9 (a larger alpha (than 0.9) did not work for some circuits in our data), while the uniqueness of the CR ensembles was maintained and the ratio of the CR ensemble neurons overlapping with RL ensembles was 26.84% (Fig. S3E), which was very similar to the case of alpha-optimized CR ensembles (Fig. 3), the size of this CR ensemble (A=0.9) was two times smaller than that of the alpha-optimized CR ensembles (Fig. S3F). Importantly, 97.82% of the neurons selected at A=0.9 were also selected in the alpha-optimized CR ensembles (Fig. S3F), suggesting that the neurons selected at the largest alpha might be more reliable and robust for the decoding among all the informative neurons. In addition, even after the removal of such “core” neurons, the remaining neurons also possessed information for the CR (Figs. S3B, D), indicating that the CR information was redundantly encoded in the dmPFC. Because this redundancy was specific to the CR ensemble and not observed in the RL ensemble, it would be interesting to investigate possible changes in this redundancy when the memory is recalled as a long-term memory (e.g. 30 days after the memory consolidation).

To evaluate the dominance of the CR ensembles vs the RL ensembles, we applied the CR decoder to predict the RL, and vice versa (Figs. 3 and S5).

### CRF models to evaluate functional connectivity

To evaluate the functional connectivity between neurons in the recorded network and the pattern completion capability of each neuron, we used conditional random fields (CRFs) as described previously (*30*), which models the conditional probability distribution of a given neuronal ensemble firing together. We used CRFs to capture the contribution of specific neurons to the overall network activity defined by population vectors belonging to a given neuronal ensemble. We generated a graphical model in which each node represents a neuron in a given ensemble and edges represent the dependencies between neurons. For training, 80% of the recorded data randomly selected from all time frames was used, and for cross-validation, the remaining 20% was used. For this analysis, binned neural activity data (1 s) were used. The model parameters were determined by the local maximum of the likelihood function in the parameter space. We constructed a CRF model in two steps: (1) structure learning, and (2) parameter learning. For the structure learning, we generated a graph structure using ℓ1-regularized neighborhood-based logistic regression(*31*). Here λs is a regularization parameter that controls the sparsity (or conversely, the density) of the constructed graph structure, leaving only relevant functional connectivity, including both coactive and suppressive relationships. A previous study showed that this number of connections was enhanced as a result of optogenetic rewiring of the local network(*31*), demonstrating the reliability of the functional connectivity estimated by CRFs models. Therefore, we also calculated the ratio of these remaining connections per all the possible connections for each neuron as a “functional connectivity” score for each node, after carefully screening the optimal λs value by maximizing the log-likelihood of the observations at the following parameter learning step. When comparing the connectivity between different ensembles (e.g. within-CR-ensemble vs within-Non-CRE) or different cell types (e.g. USR vs non-US responsive neurons), we first calculated a whole network connectivity without separating the ensembles, and further separated them into different categories. To measure which neurons were the most informative for a given stimulus (CS+ or CS−), we computed the standard ROC, taking as ground truth the timing of a particular CS. The AUC from the ROC curve that represents the performance of each neuron was calculated to compare the encoded information in different ensembles, different neuron types, and different days (e.g. before vs after the fear memory consolidation). As was recently demonstrated (*30*), high ranks for this value indicate high potential to recall the neural and cognitive representation of a given stimulus.

### Statistical Analysis

Statistical analyses in the present study were performed as described above (in “Materials and Methods” as well as in the main text and figure legends). The data that support the findings of this study are available from the corresponding author upon reasonable request. Custom codes used to analyze data in this study are available from the corresponding author upon reasonable request.

## Supplementary Materials

Supplementary Materials (Figs. S1 to S8, and caption for Movies S1 and S2) are explained in a separate document.

