## Supplementary figures for "Activity-dependent organization of prefrontal hub-networks for associative learning and signal transformation"

[1]Division of Homeostatic Development, National Institute for Physiological Sciences, 38 Nishigohnaka Myodaiji-cho, Okazaki, Aichi, 444-8585, Japan
[2]Japan Science and Technology Agency, PRESTO, 4-1-8 Honcho, Kawaguchi, Saitama, 332-0012, Japan
[3]SANKEN (The Institute of Scientific and Industrial Research), Osaka University, Mihogaoka 8-1, Ibaraki, Osaka 567-0047, Japan
[4]Division of Molecular Design, Research Center for Systems Immunology, Medical Institute of Bioregulation, Kyushu University, 3-1-1 Maidashi, Higashi-ku, Fukuoka 812-8582, Japan
[5]Department of Computer Science, Graduate School of Information Science and Technology, The University of Tokyo. 7-3-1 Hongo, Bunkyo-ku, Tokyo 113-0033, Japan.
[6]Brain Science Institute, Tamagawa University, 6-1-1 Tamagawagakuen, Machida, Tokyo, 194-8610, Japan
[7]Instituto de Neurobiologia, National Autonomous University of Mexico, Boulevard Juriquilla 3001, Juriquilla, Queretaro, CP 76230, Mexico
[8]Graduate School of Pharmaceutical Sciences, Osaka University, Yamadaoka 1-6, Suita, Osaka 565-0871, Japan
[9]United Graduate School of Child Development, Osaka University, Kanazawa University, Hamamatsu University School of Medicine, Chiba University, and University of Fukui, 2-2 Yamadaoka, Suita, Osaka 565-0871, Japan
[10]Division of Bioscience, Institute for Datability Science, Osaka University, 1-8 Yamadaoka, Suita, Osaka 565-0871, Japan
[11]Open and Transdisciplinary Research Initiatives, Osaka University, 2-1 Yamadaoka, Suita, Osaka 565-0871, Japan
[12]Graduate School of Medicine, Osaka University, 2-2 Yamadaoka, Suita, Osaka 565-0871, Japan

**This PDF file includes:**

Figures. S1 to S8
Captions for Movies S1 and S2

**Other Supplementary Materials for this manuscript include the following:**

Movies S1 and S2

Fig. S1

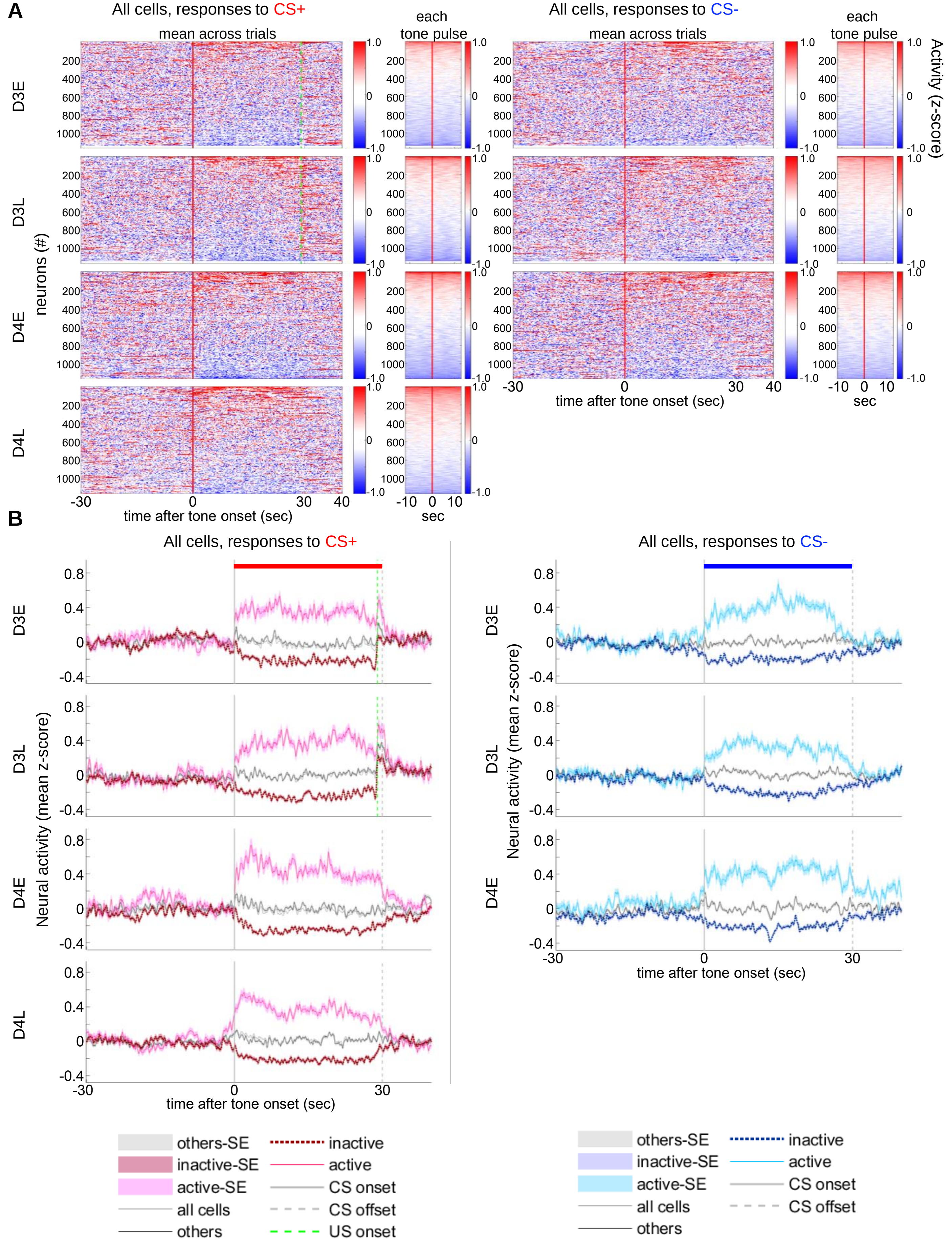
A
All cells, responses to CS+
mean across trials
each tone pulse
All cells, responses to CS-
mean across trials
each tone pulse
Activity (z-score)
D3E
D3L
D4E
D4L
neurons (#)
200
400
600
800
1000
1.0
0
-1.0
-30
0
30
40
time after tone onset (sec)
-10
0
10
sec
B
All cells, responses to CS+
All cells, responses to CS-
Neural activity (mean z-score)
0.8
0.4
0
-0.4
time after tone onset (sec)
others-SE
inactive-SE
active-SE
all cells
others
inactive
active
CS onset
CS offset
US onset

**Fig. S1. Summary of responses to CS+ and CS- of individual neurons on day (D) 3 and D4.**

To consider possible temporal changes in responses to the CSs before and after the fear conditioning, only results from mice in which neural activities were successfully recorded on both D3 and D4, from the same sets of neurons, were analyzed. (A) Mean activity over 3 CS trials, or over 87 onsets of 50-ms tone pulses during the 3 trials (D3-early[D3E]/D3-late [D3L] or D4E/D4L, respectively) for all individual neurons is plotted separately (n=1165). (B) Mean (± s.e.m.) CS responses of each category, at each temporal phase (D3E/D3L, D4E/D4L), are plotted separately.

Fig. S2

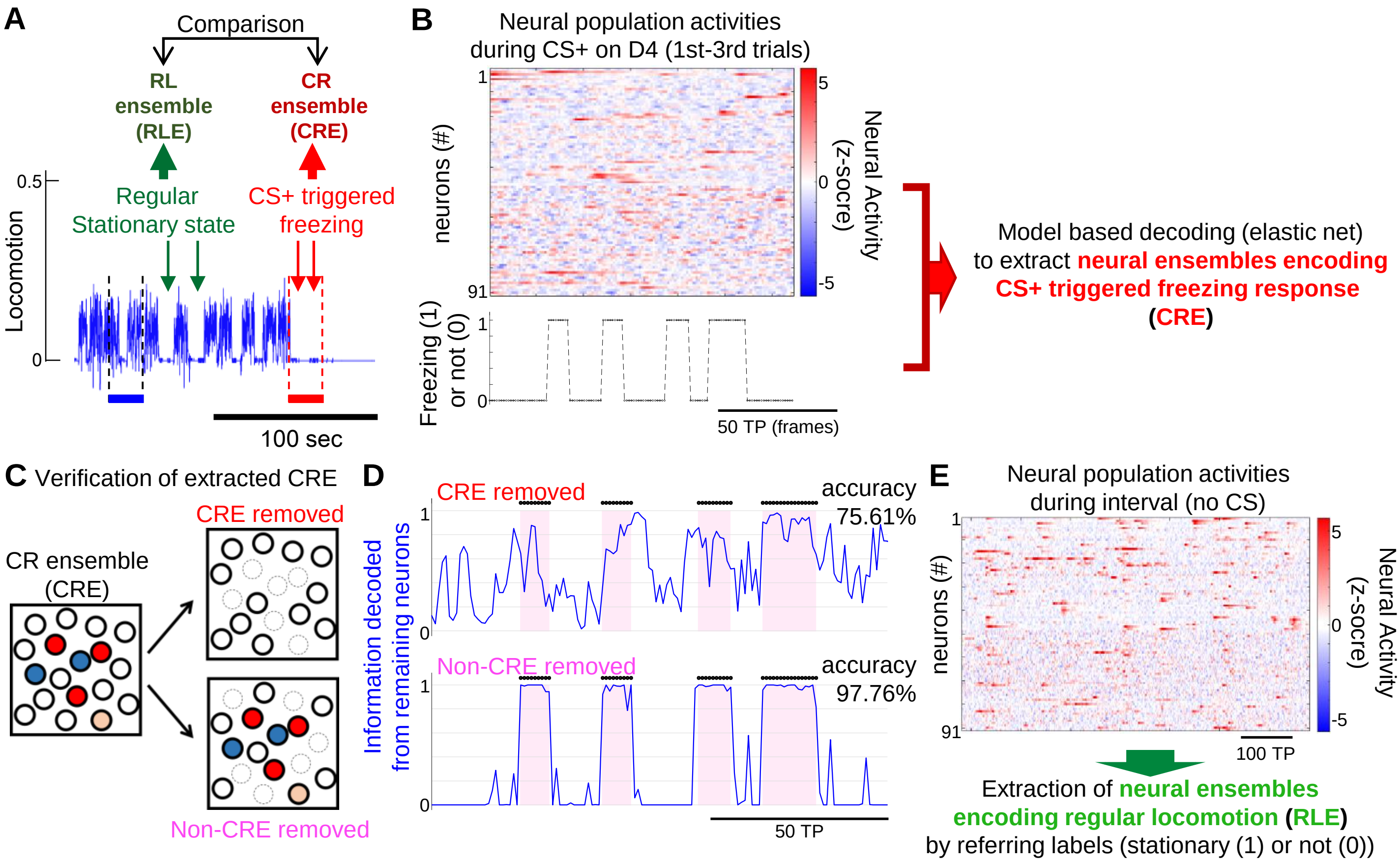


**Fig. S2. Schematic diagram showing how we extracted the CR ensemble, verified it, and compared it with the RL ensemble.**

(A) We extracted a group of neurons encoding the conditioned response (CR ensemble), and compared it with the neurons encoding regular locomotion (RL ensemble) to evaluate the uniqueness of the CR ensemble. Mice exhibited low locomotion rates during the CS+ as a CR, or during the inter-trial interval as a regular stationary state. Neurons encoding these behaviors were independently identified and compared. (B) Schematic diagram showing how we extracted the CR ensemble (CRE). The extracted CRE is shown in Fig. 2, and verified as in panel C and D. See the Materials and Methods for details. (C) The extracted CRE was verified by the comparison of the decoding performance between CRE-removed (when CRE neurons were removed) and Non-CRE removed (when non-CR-ensemble [Non-CRE] neurons were removed). The decoding performance after the removal of CRE (i.e., CRE-removed) should be substantially decreased if most of the neurons informative for the CR are selected as the CRE sufficiently. (D) Comparison of the decoding performance between CRE-removed and Non-CRE removed, revealing the poor neural information in the CRE removed. This is the result of the same mouse analyzed in Fig. 2H. Black dots on the top of the graph and pink color in the graph indicate the timing of the actual CR, while the blue line shows information encoded in this CR ensemble. (E) Schematic diagram showing how we extracted the RL ensembles. TP, time points (i.e., image frames).

### Fig. S3

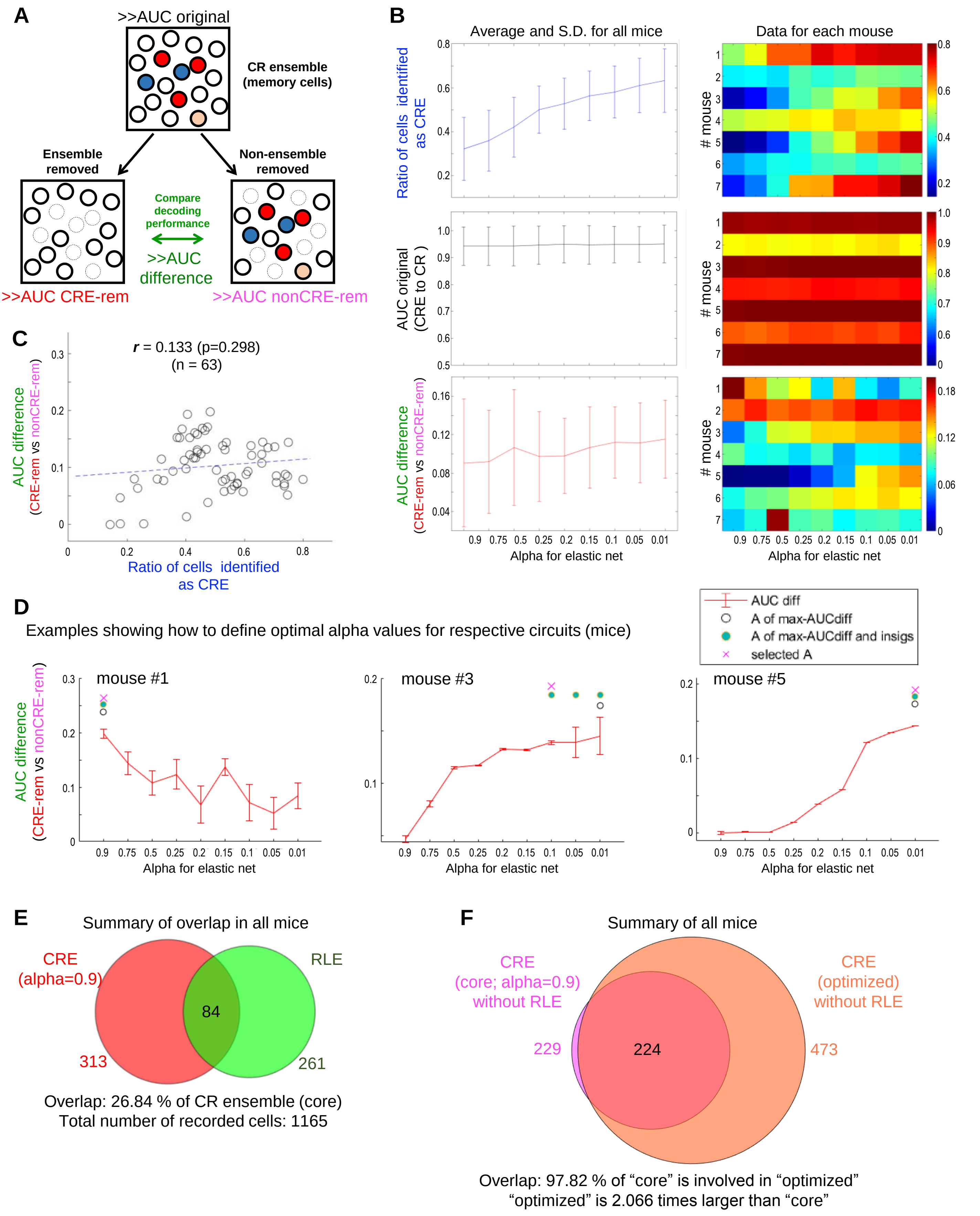

A
>>AUC original
CR ensemble
(memory cells)
Ensemble
removed
Non-ensemble
removed
Compare
decoding
performance
>>AUC
difference
>>AUC CRE-rem
>>AUC nonCRE-rem
B
Average and S.D. for all mice
Data for each mouse
Ratio of cells identified as CRE
AUC original (CRE to CR )
AUC difference (CRE-rem vs nonCRE-rem)
### mouse
Alpha for elastic net
0.9 0.75 0.5 0.25 0.2 0.15 0.1 0.05 0.01
C
r = 0.133 (p=0.298)
(n = 63)
AUC difference (CRE-rem vs nonCRE-rem)
Ratio of cells identified as CRE
D
Examples showing how to define optimal alpha values for respective circuits (mice)
AUC diff
A of max-AUCdiff
A of max-AUCdiff and insigs
selected A
mouse #1
mouse #3
mouse #5
AUC difference (CRE-rem vs nonCRE-rem)
Alpha for elastic net
E
Summary of overlap in all mice
CRE (alpha=0.9)
RLE
84
313
261
Overlap: 26.84 % of CR ensemble (core)
Total number of recorded cells: 1165
F
Summary of all mice
CRE (core; alpha=0.9) without RLE
CRE (optimized) without RLE
229
224
473
Overlap: 97.82 % of “core” is involved in “optimized”
“optimized” is 2.066 times larger than “core”

**Fig. S3. Optimization process of the size of CR ensembles by adjusting alpha values for the elastic net, which also revealed the redundant coding feature.**

(A) The AUC of the ROC was calculated to evaluate the decoding performance accurately, and to optimize the alpha value, we built a model at each alpha for each mouse (the decoding performance was calculated as “AUC original”), and further compared the difference in decoding performances between “AUC CRE-rem” and “AUC nonCRE-rem”. AUC CRE-rem is the AUC value calculated by an elastic net model built with the neurons excluding original CR ensemble neurons. AUC nonCRE-rem is the AUC value calculated by the neurons excluding neurons other than original CR ensemble neurons. The “AUC difference” between those two values was further calculated, and in principle, we defined the best alpha based on the maximum AUC difference for each mouse independently (see more details in the Materials and Methods and panel D). (B) Summary and raw data for ratio of neurons identified as CR ensemble (per whole neurons of each mouse) (top), AUC original (middle), and AUC difference (bottom), at each alpha. (C) Pearson’s correlation was used to calculate the r and p values, revealing that the ratio of neurons identified as CR ensembles (per whole neurons) and the AUC difference were not significantly correlated (n=63 samples [7 mice x 9 alphas] were analyzed to determine the possible relationship). (D) Examples showing how to determine the optimal alpha values for respective circuits (mice). In principle, we defined the best alpha based on the maximum AUC difference for each mouse independently, but in some examples as in mouse #3, several alphas revealed statistically insignificant results among the AUC differences. In this case, the largest alpha among those with the same AUC difference was selected. See more details in the Materials and Methods. (E) When alpha was fixed at alpha(A)=0.9, while the number of the neurons selected as CRE neurons were smaller than the case of the optimized alpha, the ratio of the CRE neurons that overlapped with RL ensembles was 26.84%, similar to the case of the optimized alpha as shown in Fig. 3. (F) The size of this CR ensemble (A=0.9) was two times smaller than that of the alpha-optimized CR ensembles. 97.82% of the neurons identified at A=0.9 were also selected in the alpha-optimized CR ensembles, suggesting that the neurons selected at the largest alpha 0.9 might be more reliable and robust for the decoding among all the informative neurons in the dmPFC. In addition, even after the removal of such “core” neurons, the remaining neurons also possessed information for the CR (as shown in B and D), indicating that the CR information was redundantly encoded in the dmPFC. Error bars, s.e.m.

Fig. S4

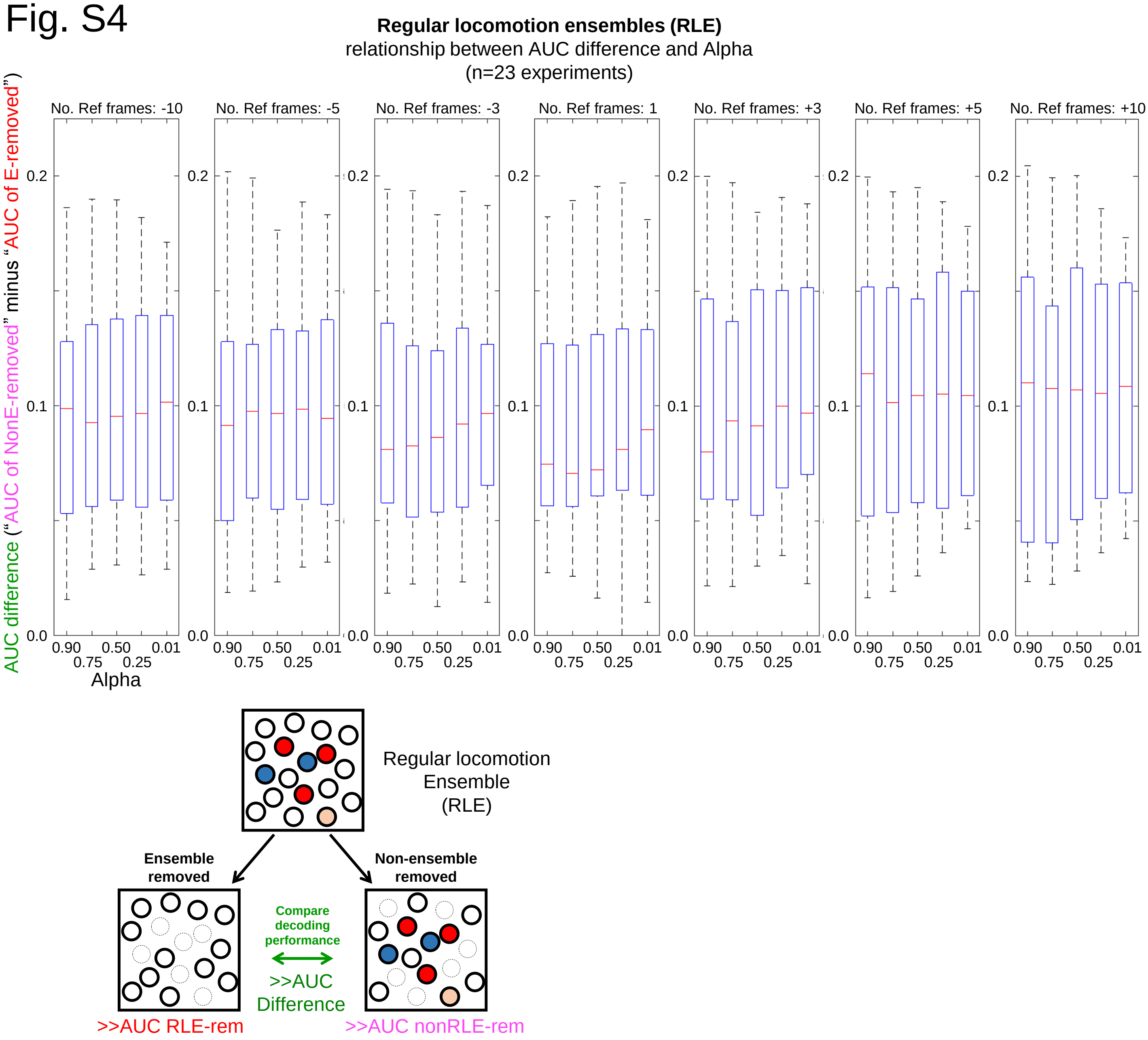


**Fig. S4. RL ensembles were not affected by the alpha of the elastic net.**

Because optimization of the alpha (hyper parameter for elastic net) was necessary to discriminate CR ensembles (Fig. S3), we also investigated the relationship between the alpha and AUC difference for RL ensembles. In addition to various alpha values, we tested various numbers of reference frames (means of the neural activities over several past or future frames were used as neural activity data to predict a single label at each time-point) to determine the potential difference in the decoding performance. We found no significant differences, however, among the various alphas, or among the different numbers of reference frames, which were evaluated by the Friedman test. Analyses shown in Fig. S5 also showed a similar independency of alpha values in the decoding performance of the RL ensembles. According to these results, we decided to fix the alpha at 0.75 to model RL ensembles, and to fix number of reference frames to one. Red bars, median; the bottom and top edges of the box indicate the 25th and 75th percentiles, respectively; whiskers extend to the most extreme data points not considered outliers (outliers were calculated by the “boxplot” function of MATLAB R2014a).

### Fig. S5

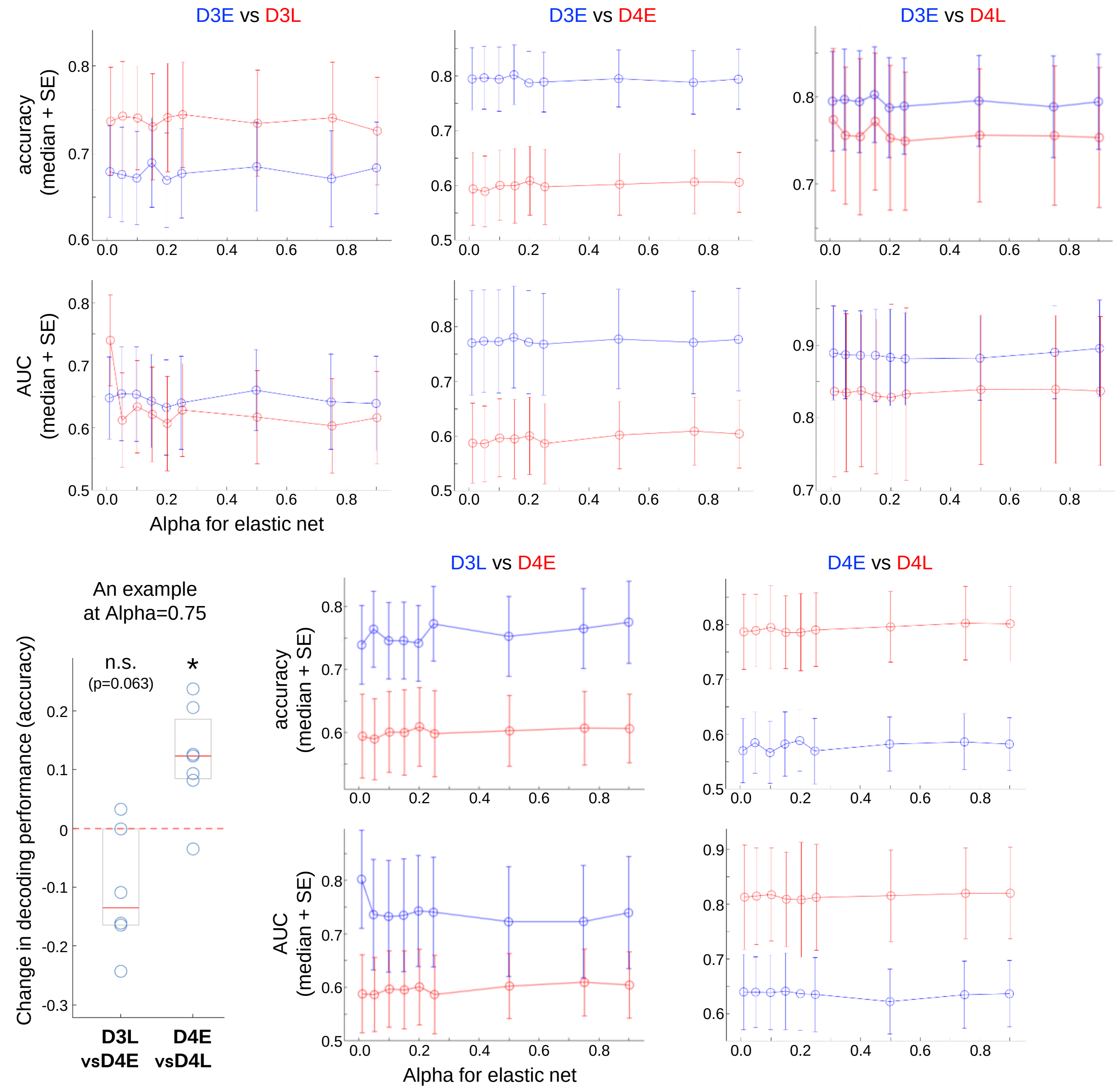


p value at each alpha (paired permutation test)

| Alpha | 0.01 | 0.05 | 0.10 | 0.15 | 0.20 | 0.25 | 0.50 | 0.75 | 0.90 |
|---|---|---|---|---|---|---|---|---|---|
| accuracy, D3EvsD3L | 0.107 | 0.219 | 0.260 | 0.113 | 0.250 | 0.309 | 0.215 | 0.244 | 0.195 |
| accuracy, D3EvsD4E | 0.000 | 0.000 | 0.000 | 0.000 | 0.000 | 0.000 | 0.000 | 0.000 | 0.000 |
| accuracy, D3EvsD4L | 0.531 | 0.500 | 0.500 | 0.531 | 0.531 | 0.594 | 0.469 | 0.531 | 0.250 |
| accuracy, D3LvsD4E | 0.156 | 0.094 | 0.156 | 0.125 | 0.219 | 0.156 | 0.125 | 0.063 | 0.031 |
| accuracy, D4EvsD4L | 0.031 | 0.016 | 0.078 | 0.016 | 0.078 | 0.031 | 0.016 | 0.016 | 0.031 |
| AUC, D3EvsD3L | 0.779 | 0.273 | 0.350 | 0.211 | 0.354 | 0.365 | 0.102 | 0.322 | 0.365 |
| AUC, D3EvsD4E | 0.219 | 0.188 | 0.219 | 0.219 | 0.250 | 0.188 | 0.188 | 0.219 | 0.156 |
| AUC, D3EvsD4L | 0.313 | 0.375 | 0.688 | 0.625 | 0.313 | 0.375 | 0.688 | 0.688 | 0.563 |
| AUC, D3LvsD4E | 0.219 | 0.563 | 0.750 | 0.656 | 0.750 | 0.563 | 0.625 | 0.563 | 0.438 |
| AUC, D4EvsD4L | 0.094 | 0.031 | 0.031 | 0.031 | 0.281 | 0.094 | 0.031 | 0.031 | 0.031 |

**Fig. S5. Summary of state-dependent change in decoding performance of RL ensembles to predict behaviors during the CS+.**

To evaluate the decoding performance, we calculated the accuracy and AUC (of the ROC) as described in the Materials and Methods. Because there was no difference among the various alphas in any estimates, as shown here and in Fig. S4, we fixed the alpha for the RL ensembles at 0.75 for further analyses. For the selected results of alpha=0.75, the data of individual circuits are also shown in the left middle panel, as in Fig. 3F. P values at each alpha (calculated by paired permutation test) are also summarized in the table.

Fig. S6

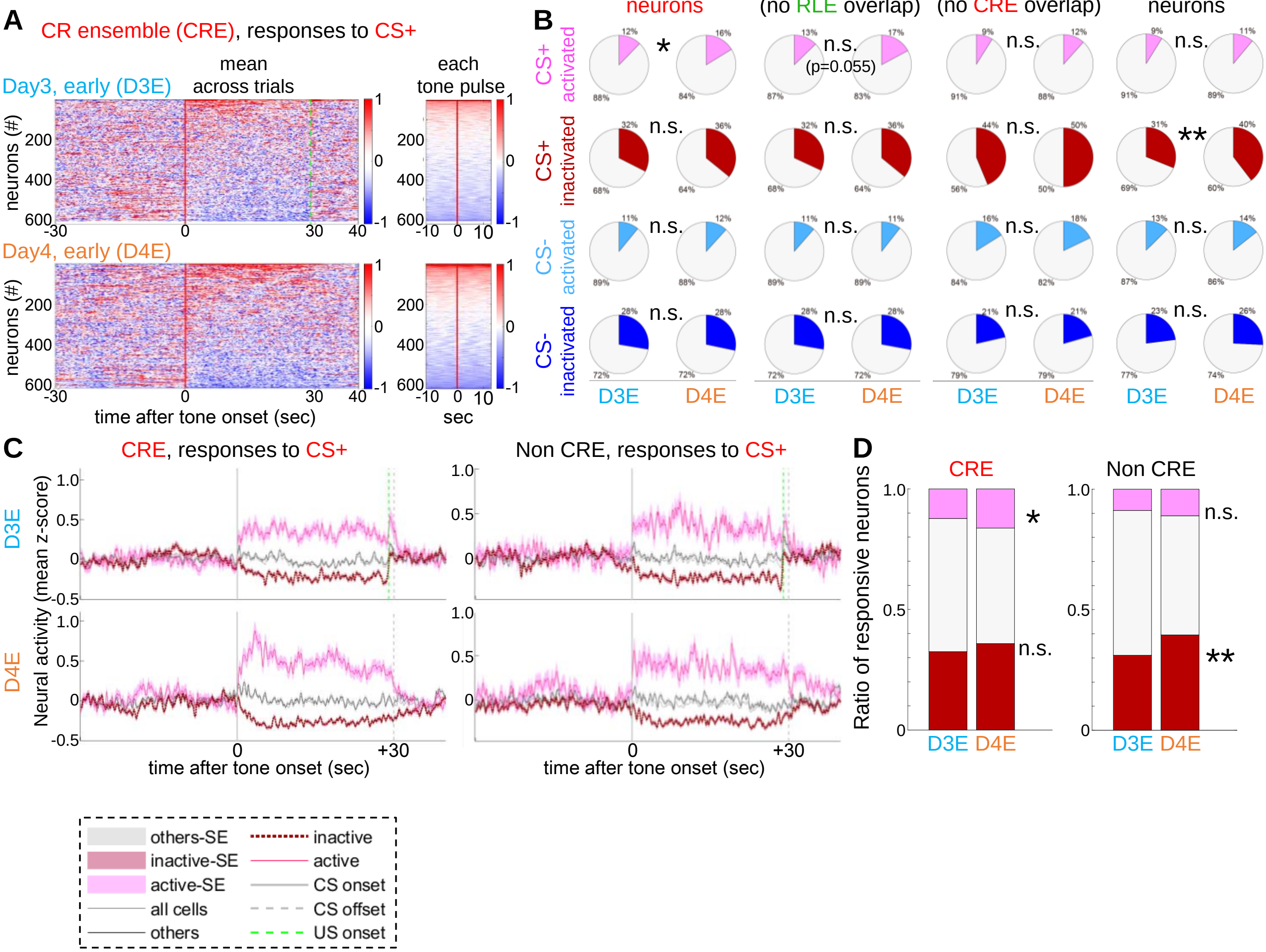


**Fig. S6. Summary of response profiles of each ensemble to CS+ and CS- on day (D) 3 and D4.**

(A) Different from Fig. S1, the mean activity over 3 CS trials, or over 87 onsets of 50-ms tone pulses during the 3 trials (D3-early [D3E] or D4E, respectively) in all individual neurons identified as CR ensemble neurons are plotted, indicating the enhanced responses at D4E compared with D3E. (B) Pie charts summarizing changes in CS responses. In CR ensembles neurons, CS+ activated neurons were slightly but significantly increased, while no change was observed for other features. On the other hand, in non-CRE neurons, only CS-inactivated neurons were significantly increased. (C) Mean CS responses (± s.e.m.) of each category in either CRE or non-CRE, at each temporal phase (D3E, D4E), are plotted separately. (D) Selected features in C were re-plotted as stacked bar graphs. A chi-square test was performed for the statistics in B and D. *p<0.05; **p<0.01; n.s., not significant.

### Fig. S7

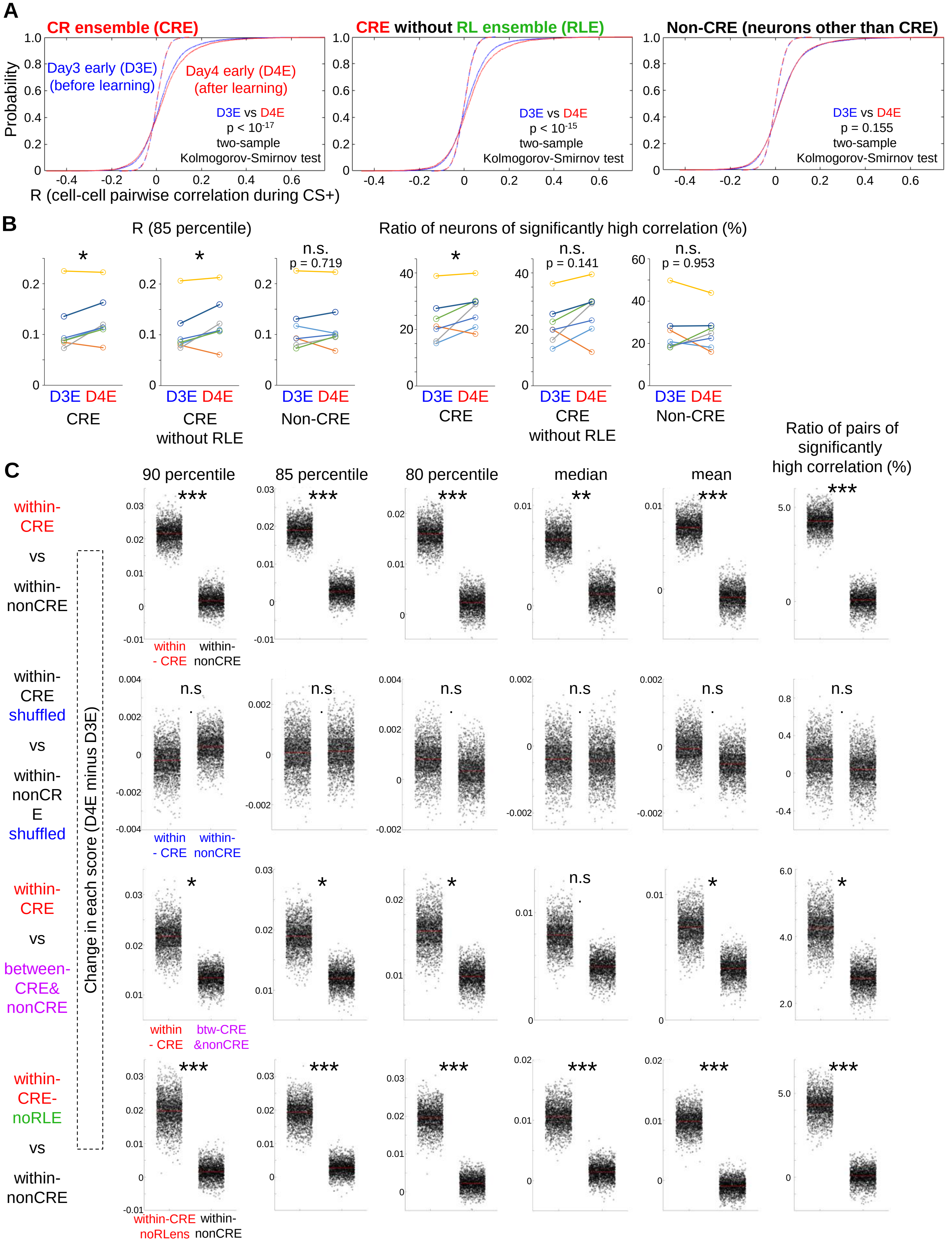

A
CR ensemble (CRE)
CRE without RL ensemble (RLE)
Non-CRE (neurons other than CRE)
Probability
Day3 early (D3E) (before learning)
Day4 early (D4E) (after learning)
D3E vs D4E
p < 10-17
p < 10-15
p = 0.155
two-sample Kolmogorov-Smirnov test
R (cell-cell pairwise correlation during CS+)
B
R (85 percentile)
Ratio of neurons of significantly high correlation (%)
n.s.
p = 0.719
p = 0.141
p = 0.953
D3E D4E
CRE
CRE without RLE
Non-CRE
C
90 percentile
85 percentile
80 percentile
median
mean
Ratio of pairs of significantly high correlation (%)
within-CRE vs within-nonCRE
within-CRE shuffled vs within-nonCRE shuffled
within-CRE vs between-CRE& nonCRE
within-CRE-noRLE vs within-nonCRE
Change in each score (D4E minus D3E)
within - CRE
within-nonCRE
btw-CRE &nonCRE
within-CRE noRLens
n.s

**Fig. S7. Change in coactivity specifically observed in CR ensembles after fear conditioning.**

(A) Cumulative curves were drawn for the data pooled by random resampling from all mice (2000 datapoints from each mouse, a total of 14,000 datapoints from 7 mice) to visualize the change in coactivity within CR ensemble neurons (CRE), CRE neurons without overlap with RL ensembles (CRE-noRLE), and neurons other than CRE (Non-CRE). Note that this random resampling is different from boot strap resampling of which results are shown in panel C (See details in the Materials and Methods). The results of the statistical comparison between day 3-early (D3E) and D4E using a two-sample Kolmogorov-Smirnov test are shown in the respective panels. Dotted lines show the results of shuffled data (no statistically significant difference in all cases). (B) Comparison between D3E and D4E for coactivity-related values was performed with the original (non-shuffled) data. In addition to the systematic analyses based on the boot strap resampling, of which results are shown in panel C, tests based on the raw data (i.e. representative values from the individual circuits) also indicated a significant enhancement of the coactivity specifically in CRE, even though the number of the samples is limited (N=7). This also demonstrated that the results shown in the panel A and C did not derive from artificially enhanced marginal differences. (C) Detailed and systematic analyses of the change in coactivity in dmPFC circuits after the fear conditioning. Changes in the correlation coefficients R (day 4-early [D4E] minus D3E) of all chronically observed circuits (N=7 mice) are plotted as a result of bootstrap resampling (2000 times) performed to systematically compare all the raw R data of whole mice, for those of CR ensemble neurons (within-CRE) or for those other than CR ensemble neurons (within-nonCRE). Because the panel A indicates that the clear change in R between D3E and D4E was observed only for positive correlations specifically in CRE, we statistically tested changes in the 90th, 85th, 80th, and 50th percentiles, mean, and ratio of pairs of the significantly high correlation for each category. We also revealed results for shuffled data and for within-CRE vs between-CRE&nonCRE (coactivity between CRE neurons and Non-CRE neurons), suggesting that enhanced coactivity within the CRE after the fear conditioning was specific. Results of CRE-noRLE were also consistent with those of CRE. A paired permutation test was used for the statistics in B. The data obtained by bootstrap resampling shown in C were statistically analyzed as described in the Materials and Methods. *$p<0.05$; **$p<0.01$; ***$p<0.001$; ****$p<0.0001$; n.s., not significant. Red bars, median; gray boxes in panel C indicate the 25th and 75th percentiles.

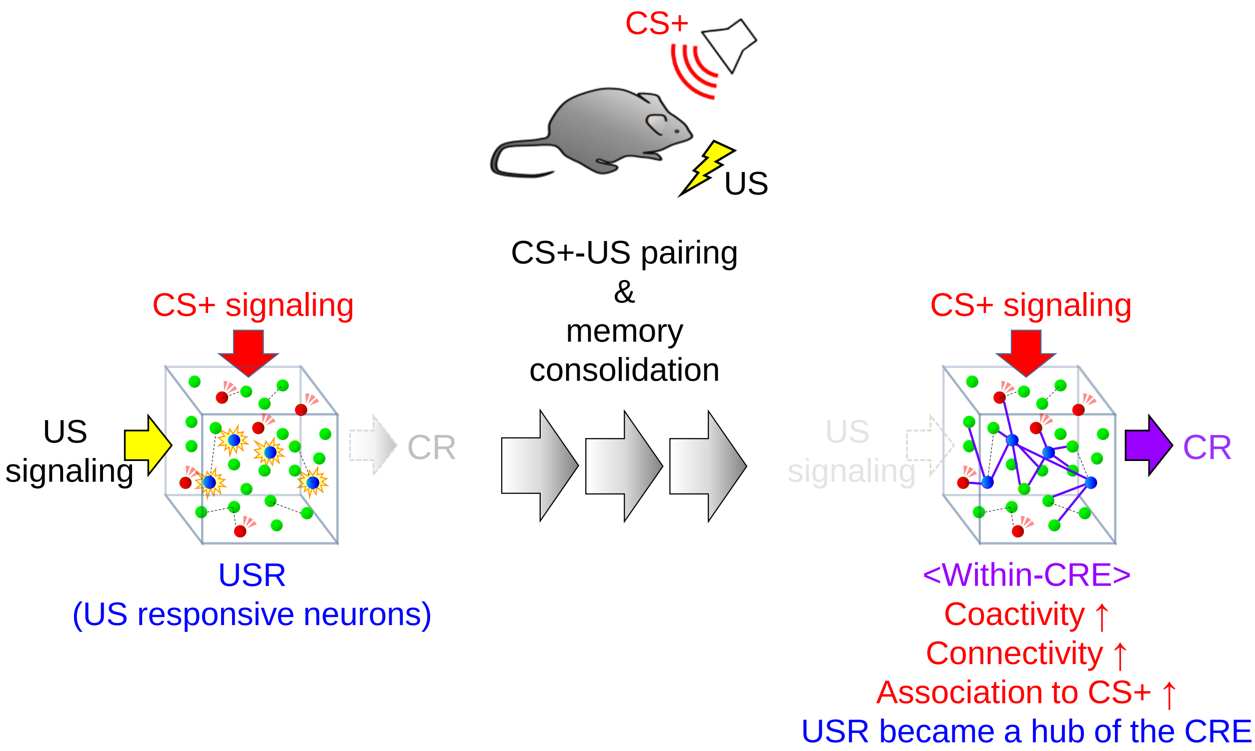


**Fig. S8. Summary of the present study.**

We demonstrated that the repeated CS+-US pairing for the associative learning drives the dmPFC reorganization to generate novel and unique neural circuits for CS-to-CR transformation, with enhanced internal coactivity, connectivity, and association with the CS+. Upon this prefrontal reorganization to encode associative memory, neurons activated by the US during fear conditioning, which were originally less associated with the CS+ network, were anterogradely and predominantly integrated into the CR ensemble (CRE) with enhanced association with the CS+ network. The eventual network stemming from these USR gained typical features of pattern completion cells of the CRE, which are supposed to work as a hub in the prefrontal networks to predominantly relay the CS+ information and promote the CR.

**Movie S1.**

An example of spontaneous activities in dmPFC on day4, detected by changes in GCaMP6f signals in a field of view. (left) original GCaMP6f signal, (middle) baseline subtracted signal shown in magenta, (right) baseline subtracted signal is shown in magenta over a background of the baseline-image shown in gray.

**Movie S2.**

An example of GCaMP6f signals on day3, during fear conditioning. Baseline subtracted signal is shown in magenta, and merged over the baseline-image shown in gray. The timings of the CS and US presentation were indicated at the left upper corner in the movie. The CS and US presentation did not disturb the image acquisition.
